# Data-driven analyses of social complexity in bees reveal phenotypic diversification following a major evolutionary transition

**DOI:** 10.1101/2024.02.09.579609

**Authors:** Ohad Peled, Gili Greenbaum, Guy Bloch

**Affiliations:** Department of Ecology, Evolution, and Behavior, The Silberman Institute of Life Sciences, The Hebrew University of Jerusalem; Jerusalem, Israel

## Abstract

How social complexity evolved is a long-standing enigma. In most animal groups, social complexity is typically classified into a few discrete classes. This approach is oversimplified and constrains our inference of social evolution to a narrow trajectory consisting of transitions between classes. This approach also limits quantitative studies on the molecular and environmental drivers of social complexity. However, the recent accumulation of relevant quantitative data has now set the stage to overcome these limitations. Here, we propose a data-driven approach for studying the full diversity of social phenotypes. We curated and analyzed a comprehensive dataset encompassing 17 social traits for 77 species and studied the evolution of social complexity in bees. We found that corbiculate bees — honey bees, stingless bees, and bumble bees — underwent a major evolutionary transition ∼70 mya, which is inconsistent with the stepwise progression of the social ladder conceptual framework. This major evolutionary transition was followed by a phase of substantial phenotypic diversification of social complexity. Non-corbiculate bee lineages display a continuum of social complexity, ranging from solitary to simple societies, but do not reach levels of social complexity comparable to those of corbiculate bees. Bee evolution provides a unique demonstration of a macroevolutionary process in which a major transition removed biological constraints and opened novel evolutionary opportunities, driving the exploration of the landscape of social phenotypes. Our approach can be extended to incorporate additional data types and readily applied to illuminate the evolution of social complexity in additional animal groups.

## Introduction

One of the most fascinating examples of increased complexity in biological systems is the evolution of animal societies. Sociality has evolved multiple times across independent taxonomic groups and has played a key role in the evolution of animals^1–4^. Despite remarkable variation among animal societies^4,5^, comparative studies have typically classified levels of social complexity based on few qualitative traits, such as group composition, reproductive skew, and parental care (e.g., in primates^6–9;^ in birds^10,11^; in insects^12–15)^. This approach was crucial for the development of influential theories such as kin selection^16^, major transitions in evolution^17^, and developmental plasticity^18^. However, focusing on a few qualitative traits forces social phenotypes to fit into a small number of coarsely defined classes (e.g., pair-living, subsocial, eusocial) assumed to represent a set of evolutionary transitions^19–23^. This limited set of transitions imposes a narrow trajectory for the evolution of social complexity^19^. Such qualitative classifications fail to capture the full diversity of social phenotypes within these classes. For example, all ants and termites are typically classified as advanced eusocial despite including species with simple colonies of a few dozen individuals to complex societies composed of millions of individuals^15,24^. This approach may mask important evolutionary processes such as phenotypic diversification.

Insects from the order Hymenoptera provide a powerful system with which to study the evolution of social complexity because species within the same taxonomic lineage often exhibit diverse social phenotypes^12,25,26^. Theories on the evolution of social complexity in insects can be divided into two main frameworks. The first theory emphasizes a single pivotal increase in the level of social complexity that leads to an irreversible point-of-no-return, known as superorganismality^13,17,21,27,28^. This framework focuses on caste differentiation processes producing a highly fertile caste (queens or kings) which functions as the reproductive entity of the superorganism, and workers with severely reduced reproductive potential, which are functionally analogous to somatic cells in a multicellular organism. The second framework, commonly termed ‘the social ladder’, assumes a stepwise evolutionary trajectory passing through a few key levels of sociality^12,15,29^. The less complex societies are viewed as primitive states, metaphorically comparable to lower rungs on a ladder^29–31^. Despite the long tradition and significant contribution of the social ladder framework, it remains debated whether sociality in insects evolved along similar evolutionary trajectories across different taxonomic lineages^19,20,23,32–34^.

Both theories have suffered from repeated semantic disagreements regarding their definitions^20,22,23,34^, a discussion that hampered the use of comparative methods to identify factors associated with variation in social complexity^19,20,26^. Indeed, recent attempts to understand the molecular underpinnings of the evolution of social complexity have had limited success^29,35–40^. This lack of substantial progress can be partially attributed to the limited inference that can be made when using a small number of traits as proxies for social complexity^19,20^, given that traits contributing to complex phenotypes may be shaped by various selection pressures, ecological niches, preadaptations, pleiotropic effects, and developmental constraints^41–46^. Previous attempts to apply quantitative indices for social complexity focused on few traits and were restricted by both the number of species included and the range of taxonomic sampling^20,47–50^, limiting their capacity to identify and characterize macroevolutionary processes.

The increased availability of life history, behavioral, morphological, and molecular data for related species differing in social phenotypes, coupled with well-resolved phylogenetics and appropriate computational methods, sets the stage for a detailed investigation of social complexity phenotypes^51–58^. Here, we adopt a data-driven approach to untangle the evolutionary history of social complexity in bees, which does not assume that there are specific social classes or predetermined evolutionary trajectories. Such a model-free approach enables the analysis of complex interactions among traits in a high-dimensional space^59–62^, and can potentially resolve controversies arising from a qualitative representation of social complexity.

## Results

We generated a quantitative dataset consisting of 17 traits related to social complexity for 77 bee species exhibiting diverse levels of social complexity (Fig. 1; see Fig. S1 for species names). Through an extensive literature survey, we identified reliable quantitative social traits that are comparable across species (Fig. 1a and Table S1). Our dataset includes colony-level traits (e.g., colony size, colony longevity, colony reproductive skew, type of brood care), life-history traits (e.g., overlap of generations, colony founding), and morphological traits (e.g., queen-worker and worker-worker size polymorphism). We did not include traits that cannot be properly compared across species (e.g., behavioral repertoire) or that are not available for many taxa (e.g., detailed morphological and anatomical traits). The 77 bee species in our dataset were selected based on the availability and quality of published information, aiming to accurately represent the broad spectrum of social complexity and phylogeny (Fig. 1b). Some clades are characterized by a single level of social complexity (e.g., honey bees, stingless bees, bumble bees, Megachilidae solitary bees), while other clades encompass multiple levels of social complexity (e.g., sweat bees, orchid bees, and Xylocopinae bees; Fig. 1b).

**Figure 1.**
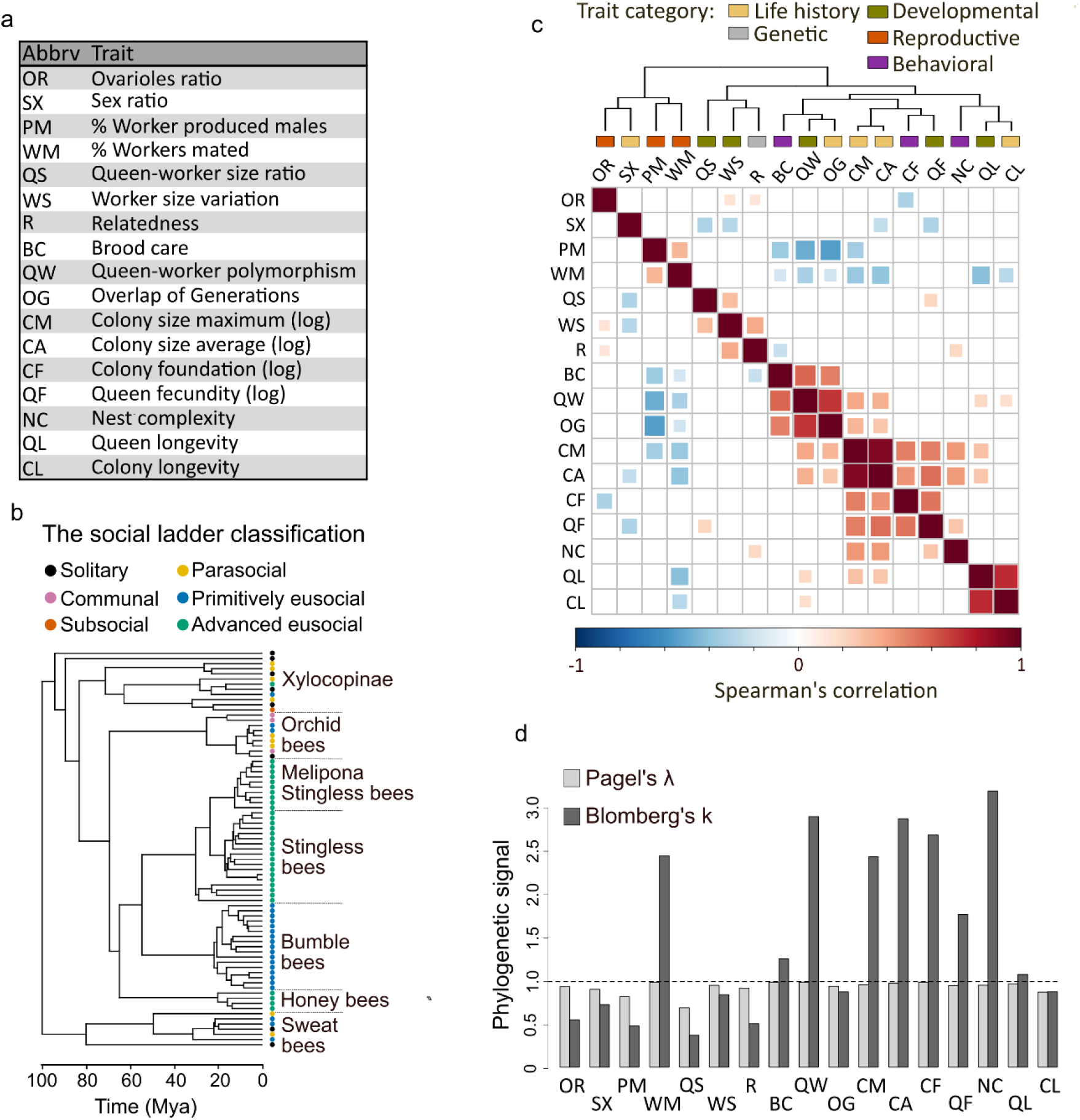
Overview of our dataset consisting of 17 traits and 77 bee species. **(a)** The list of social complexity related traits in our dataset. **(b)** Time-calibrated phylogenetic tree of the species in our dataset. See Fig. S1 for species names. Colors correspond to the common social ladder classification of Michener^12^. **(c)** A heatmap summarizing the correlation between social complexity traits in our dataset. The color and size of the boxes correspond to the strength of the Spearman correlation coefficient, with positive correlations in red and negative in blue. Only statistically significant correlations are shown (*p*-value<0.05 after FDR correction). Hierarchical clustering is shown on top of the correlation matrix. The color bars on top of the matrix classify each trait into one of five broadly defined categories. (**d**) The phylogenetic signals for each social complexity trait in our dataset; Pagel’s λ are in dark gray and Blomberg’s K are in light gray. The dashed horizontal line delineates a neutral Brownian motion process. All traits showed a significant phylogenetic signal (*p*-value<0.001 for both measures after FDR correction) when compared to a null model in which traits evolve independently of phylogeny.

Many traits in our dataset, including colony size-related traits (CA, CM, CF), queen fecundity (QF), and nest complexity (NC) are highly correlated (Fig. 1c). Interestingly, the defining attributes of eusociality^15^ – brood care (BC), overlap of generations (OG), and queen-worker polymorphism (QW) – were strongly correlated with each other, but not with many of the other social traits in our dataset. Likewise, colony and queen longevity (CL, QL) are correlated with each other but not with other traits. Relatedness (R) between females, a putative driver for social evolution, was not correlated with most traits. The percentage of mated workers (WM) was negatively correlated with many other social traits. Overall, the correlation matrix suggests that not all social traits are necessarily positively correlated and that multiple combinations of traits may contribute differently to an increase in social complexity.

Phylogeny significantly accounted for variation in social complexity. There is a statistically significant phylogenetic signal (*p*<0.001) for all traits (Fig. 1d) using either Pagel’s lambda or Blomberg’s K. The two traits with the lowest phylogenetic signal were worker-produced males (PM) and queen-worker size ratio (QS). For seven traits, Blomberg’s K values were >1: colony size-related traits, queen-worker polymorphism, and the percentage of mated workers, which is consistent with higher-than-expected similarity based on phylogenetic relatedness alone. These traits are also those that were highly correlated. Nest complexity showed the highest Blomberg’s K value, probably due to its high score in stingless bees, which build remarkably elaborate nests. The strong phylogenetic signals for the social traits are consistent with the occurrence of similarly high levels of social complexity in honey bees, stingless bees, and bumble bees which are represented by a relatively large number of species in our database (Fig. 1b).

### The phenotypic space of social complexity in bees

We used dimensionality reduction analyses to describe the phenotypic space of social complexity using the 17 traits in our dataset. We employed both principal component analysis (PCA) and uniform manifold approximation and projection (UMAP). These analyses are complementary; PCA describes the global structure of the phenotypes of the species in our dataset, whereas UMAP emphasizes within-group variation, which can add fine resolution to clustering in the phenotypic space. Given the significant phylogenetic signals for all traits in our dataset (Fig. 1d), we applied a phylogenetic correction for all downstream analyses. The clustering in UMAP was consistent over multiple iterations and parameters (Fig. S2), indicating that the overall structure of the data we obtained is robust regarding the relations between species. In the PCA, the broken stick method^63^ suggested four potentially informative PCs which together explain ∼70% of the variation. PC1 and PC2 accounted for 28% and 15% of the variation, respectively (Fig. 2a; data on all four main principal components are shown in Figs. S3 and S4). Both the PCA and UMAP identify clusters of species occupying clear areas in the phenotypic space, but the composition and structure of the clusters were partially different across analyses (Fig. 2a).

**Figure 2.**
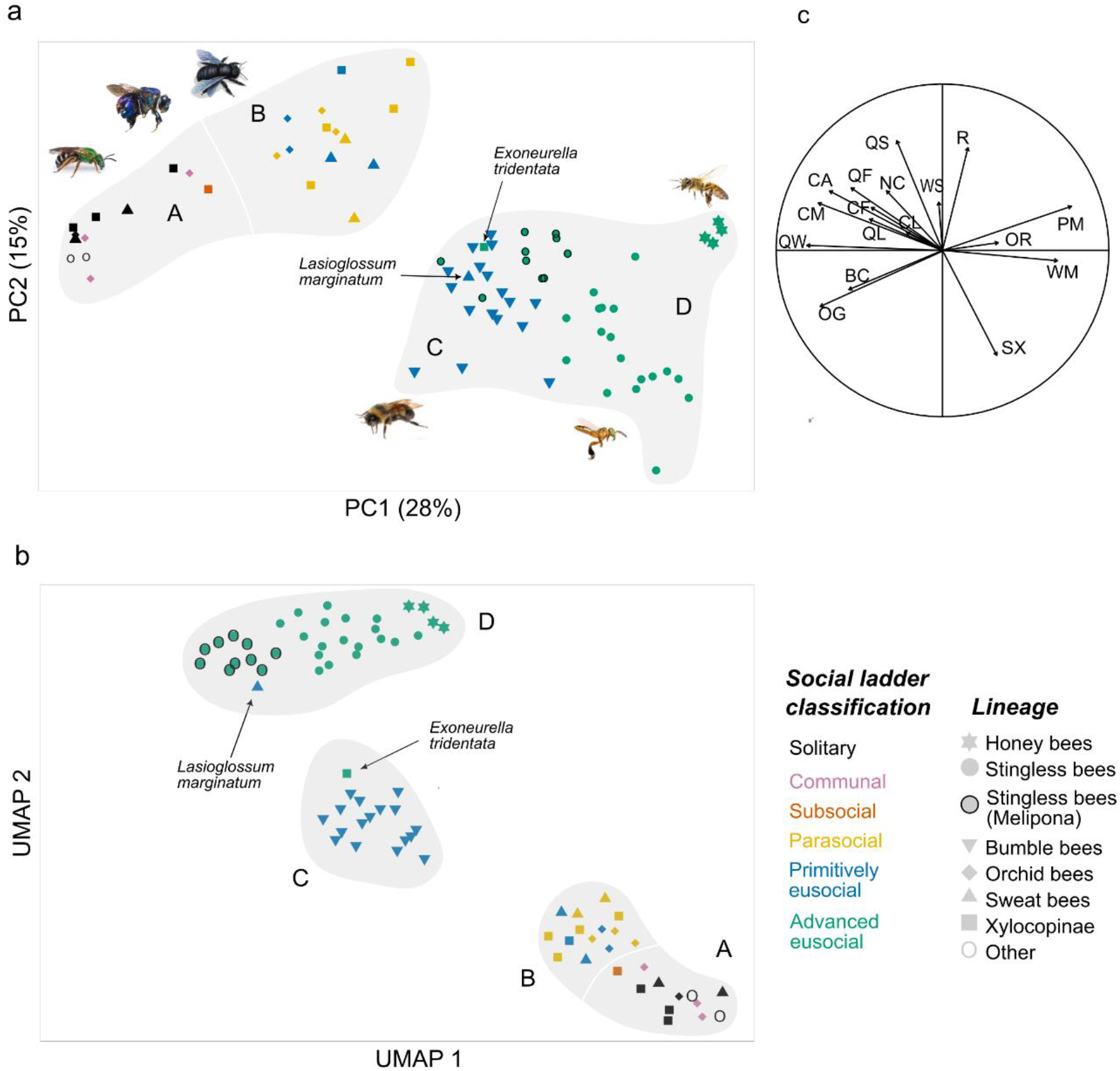
The phenotypic space of social complexity in bees. (a) Phylogenetically corrected Principal Component Analysis (PCA). The plot shows Principal Components 1 and 2 (PC1 and PC2), which together account for 43% of the variation in our dataset. Symbol color corresponds to the commonly used social classification and its shape to the species taxonomic clade (legend on the bottom right). *Habropoda labriosa* and *Megachile rotundata* are not included in the main clades in our database and were therefore marked as ‘other’. (b) Phylogenetically corrected Uniform Manifold Approximation and Projection (UMAP). We visually identified four putative phenotypic groups, marked with capital letters A to D. (c) The variable correlation plot describes the contribution of traits to PC1 and PC2, with label abbreviations detailed in Fig. 1a. The direction of each vector corresponds to the PCA in (a) and indicates the proportion of its contribution to PC1 and PC2, and its length indicates the amount of contribution.

In UMAP (Figs. 2b and S2), clustering suggested a clear separation between (i) honey bees and stingless bees (ii) bumble bees, and (iii) the remaining species. In the PCA plots (Fig. 2a and S4), bumble bees and stingless bees overlapped in PC1 and PC2 but not in PC3 and PC4, whereas honey bees were clustered separately. In addition, solitary, communal, and subsocial species were clustered separately from parasocial (often termed facultatively social^64,65^) and non-*Bombus* primitively eusocial species in most of the PC combinations. Considering both PCA and UMAP, we identified four main phenotypic groups that occupy well-defined areas in the phenotypic space representing different social complexity phenotypes (hereafter groups; grey shaded areas labeled A-D in Fig. 2a-b). The main groups correspond to: (A) solitary, communal, and subsocial species; (B) parasocial and non-*Bombus* primitively eusocial species; (C) bumble bees; (D) honey bees and stingless bees. The clearest and widest separation in both the UMAP and PCA is between the monophyletic group of corbiculate bees — honey bees, stingless bees, and bumble bees (groups C and D; hereafter corbiculate bees) — to the rest of the species (groups A and B; Fig. 2a-b). This partition reflects the separation of species with obligate sociality (C and D) from species with facultative or no social lifestyle (A and B). Notably, facultative and obligate sociality was not one of the traits included in our dataset, indicating that our data-driven approach successfully captured key attributes of social complexity that were not explicitly defined.

The clusters in our phenotypic space differ from the commonly used social ladder classification (color-coded in the phenotypic spaces in Figs 2, S2, S4). Bees assigned to different qualitative levels of sociality clustered together in our phenotypic space. The change from solitary lifestyle (group A) to parasocial and non-*Bombus* primitively eusocial species (group B) was gradual rather than stepwise as predicted by social ladder models. Another apparent disagreement with social ladder models is that species that are classified as primitively eusocial do not occupy a defined phenotypic area, but rather are separated into two different clusters: one that includes bumble bees, and one that includes the other primitively eusocial species as well as some species that are typically classified as parasocial. *Exonourella tridentata* (Allodapini, typically classified as advanced eusocial) and *Lasioglossum marginatum* (Halictidae; typically classified as primitively eusocial) were the only species in groups C and D that do not belong to the corbiculate bees. These two species were associated with different clusters across analyses (Fig. S4 and S5). Notably, *Bombus atratus* (marked in Fig. 2a) is positioned in the margins of the bumble bee cluster, most likely attributed to its perennial colonies, which can be larger than those of some of the stingless bees.

In the PCA, the species clusters are not tight, with each occupying a large area in the phenotypic space. Notably, the within-group variation is as large as the between-group variation (i.e., the size of each group in the PCA is as large as the distances between the groups), highlighting significant variation in social complexity that is entirely overlooked in the commonly used qualitative classifications. An important example is the large phenotypic space occupied by advanced eusocial species, suggesting that species typically assigned to this class vary substantially in social traits (group D in Figs. 2 and S4). We computed the convex hulls of regions in the phenotypic space occupied by corbiculate bees and the remaining species, considering all four main PCs, and found that the area occupied by corbiculate bees is twice as large (see Fig. S6 for visualization of the analysis for PCs 1-2). Our analyses suggest that group D can be further partitioned into three subgroups: honey bees, *Melipona* genus stingless bees, and non-*Melipona* stingless bees (Figs. 2a-b). The bumble bee group was consistently partitioned along the common distinction between mass provisioners (pocket-making) and progressive provisioners (pollen-storers; shown in UMAP analysis Fig. S2). Analyzing the correlation of traits with the PCs (Fig. 2c) suggests that the variation characterizing the corbiculate bees is attributed to most of the traits in our dataset, whereas variation in groups A and B (e.g., sweat bees, orchid bees, and Xylocopinae bees) stemmed primarily from variability in brood care (BC), the percentage of worker-produced males (PM), overlapping generations (OG), sex ratio (SX), and relatedness (R).

### The evolutionary history of social complexity in bees

We conducted ancestral state reconstruction using principal component values as a quantitative estimate for the social complexity phenotype to gain insight into the evolutionary history of social complexity (Fig. 3a and S7). To visualize the evolutionary reconstruction in more than a single dimension, we projected the ancestral state reconstruction into different combinations of the four main PCs (Figs. 3b and S8). These analyses revealed evolutionary flexibility and repeated changes in the social complexity phenotype of species from groups A+B along PCs 1-3. The reconstructed evolutionary trajectory leading to the corbiculate bees showed a relatively consistent increase in social complexity values along PC1 over time (Fig. 3a). We identified a presumed point-of-no-return that is consistent with an irreversible phenotypic threshold in PC1 and was crossed by the corbiculate bees, as well as by *E. tridentata* and *L. marginatum* (dashed line in Fig. 3a). The evolutionary reconstruction of PC2, with high variation contributed by the traits Q-W body size ratio, sex ratio, and relatedness, also shows an early separation of the corbiculate bees from the other species. The social complexity of bumble bees, stingless bees, and honey bees later diversified within the phenotypic space of PC1 and PC2. For example, the stingless bees diverged from the rest of the stingless bees to occupy a distinct phenotypic space (Fig. 3b). The corbiculate bees show the largest divergence during the reconstruction of PC3 and PC4 (Fig. S7 and S8). Our evolutionary reconstructions suggest that the high social complexity of *E. tridentata* and *L. marginatum* reflects a later increase in social complexity through different evolutionary routes than that which led to high social complexity in corbiculate bees (Figs. 3b, S7).

**Figure 3.**
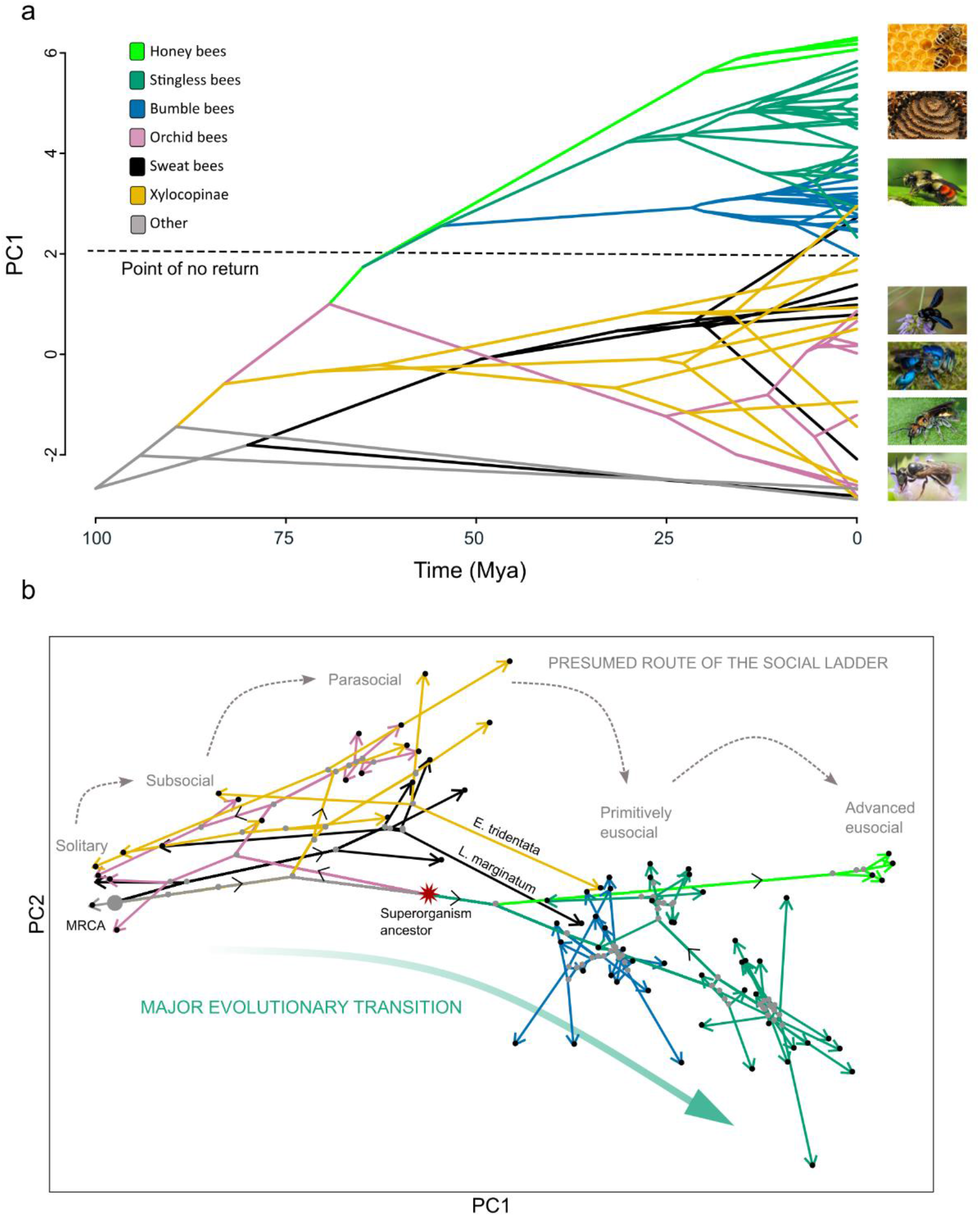
Ancestral state reconstruction for the evolution of social complexity. Values were obtained from the PCA (Fig. 2a). **(a)** A phenogram of the evolutionary trajectory of PC1 over time. The root was set to a minimum value based on the solitary *M. rotundata*. Colors represent the main taxonomic lineages in our dataset, broadly aligning with the color scheme in Fig. 2. A dashed line indicates a putative point-of-no-return beyond which species cannot revert to lower levels of social complexity. **(b)** Integration of the PCA phenotypic space with ancestral state reconstruction analysis of PC1 and PC2. Arrows depict the inferred evolutionary route of species within the phenotypic space. The arrowheads and the black dots mark the position of extant species in our dataset. Gray dots denote the assumed social phenotype of putative ancestral species (i.e., internal nodes in the phylogeny). Large gray dot depicts the most recent common ancestor of all species (MRCA). The red star marks the putative common ancestor of corbiculate bees. Gray dashed arrows illustrate a stepwise increase in social complexity as predicted by social ladder models. The thick green arrow suggests a putative trajectory consistent with a major transition in social complexity. Similar analyses for PCs 1-4 are presented in Fig. S8).

A shift-detection analysis using the values of PC1 and PC2 revealed a single major evolutionary shift, which occurred in the ancestor of corbiculate bees (green dot in Fig. 4); the same result was obtained by directly analyzing the traits instead of PCs, Fig. S9b). Models assuming a larger number of shifts, as hypothesized, for example, by social ladder models, received significantly less support (Fig. S10). We did find additional phenotypic shifts when including all four main PCs, but these were in the specific branches leading to honey bees, stingless bees, *Melipona* genus, and bumble bees (fig. S9a). This suggests that PC3 and PC4 offer a more detailed description of phenotypic changes in these taxonomic lineages. Sensitivity analyses suggested that the evolutionary trajectories, as well as the phenotypic shift we identified, are not biased by the taxonomy representation, missing data, or specific traits in our dataset (Figs. S11-S13). We, therefore, conclude that after the corbiculate bees diverged from the remaining species, they crossed a major evolutionary transition in social complexity from which a return to a simpler social organization is not likely.

**Figure 4.**
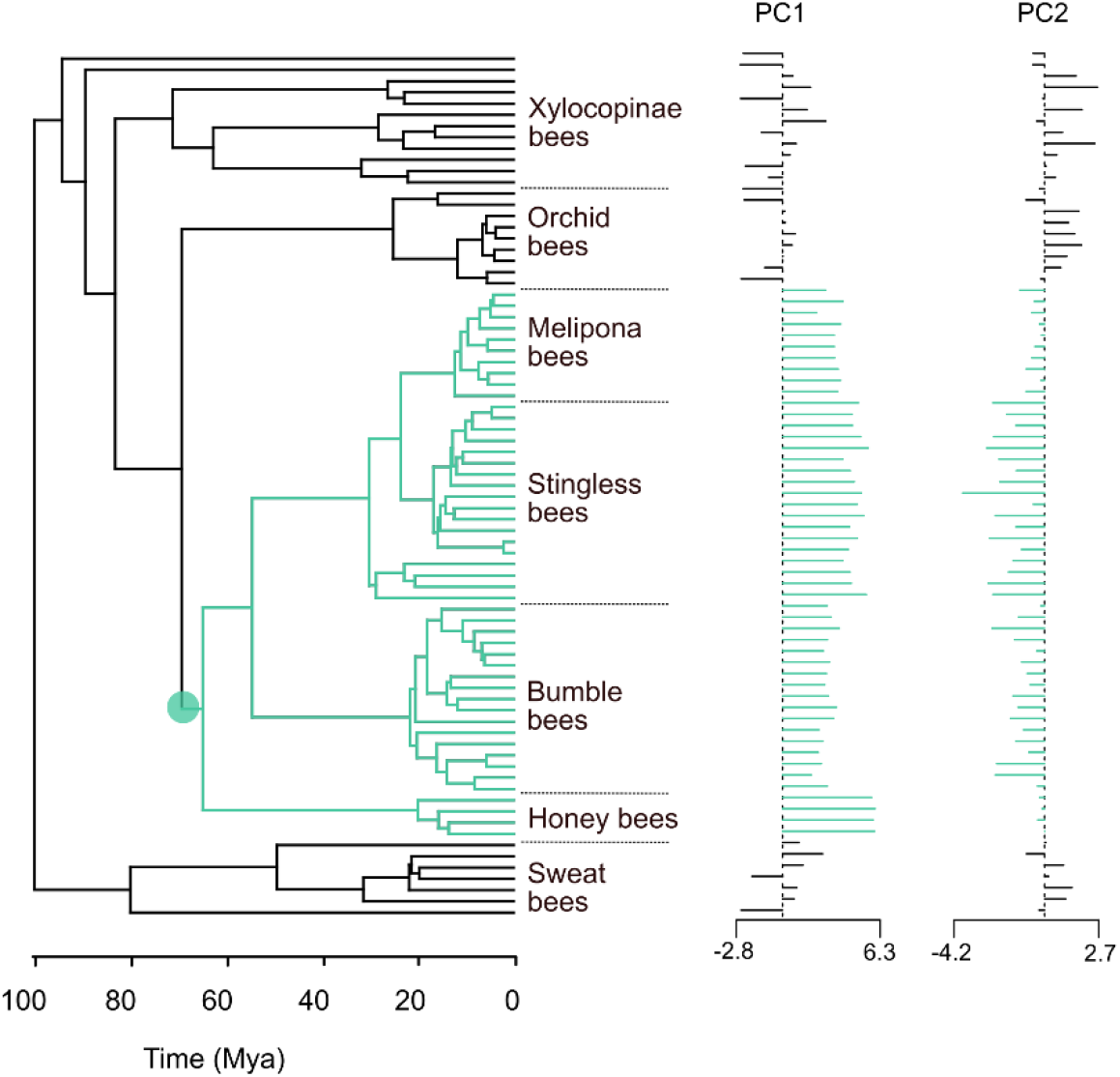
Shift detection analysis. On the left, a single shift detected on the phylogenetic tree. The green circle indicates the inferred position of adaptive regime shift from PC1 and PC2, marking the ancestor of corbiculate bees. On the right, bars show the PC1 and PC2 values for each species. Analysis for PCs 1-4 is shown in Fig. S9a.

### Evolutionary transition leading to phenotypic diversification

To quantitatively study the directionality of the evolutionary trajectories and phenotypic diversifications, we analyzed the angle of change and the size of each step in the phenotypic space in the reconstructed trajectories (illustrated in Fig. 5a). We used the trajectories in the phenotypic space of PCs 1-4. Before the divergence of the corbiculate bees, we observe a high variance of the degree distribution for all species (Fig. 5b). However, the trajectory of corbiculate bees after the divergence showed a statistically significant lower variance of the degree distribution compared with the rest of the species (*F*-test, 7.31, *p*<0.001), which was seven times higher. We found a higher and closer to zero mean angle in the reconstructed evolutionary trajectory of the corbiculate bees compared with species of other clades after their divergence (3.01±0.2 and 2.5±0.54 radians for corbiculate and other species, respectively; *t*-test, *p*<0.001) These findings are consistent with the hypothesis that the increase in social complexity over time was more directional and less flexible in the corbiculate bees compared to the branches leading to the sweat bees, orchid bees, and Xylocopinae species.

**Figure 5.**
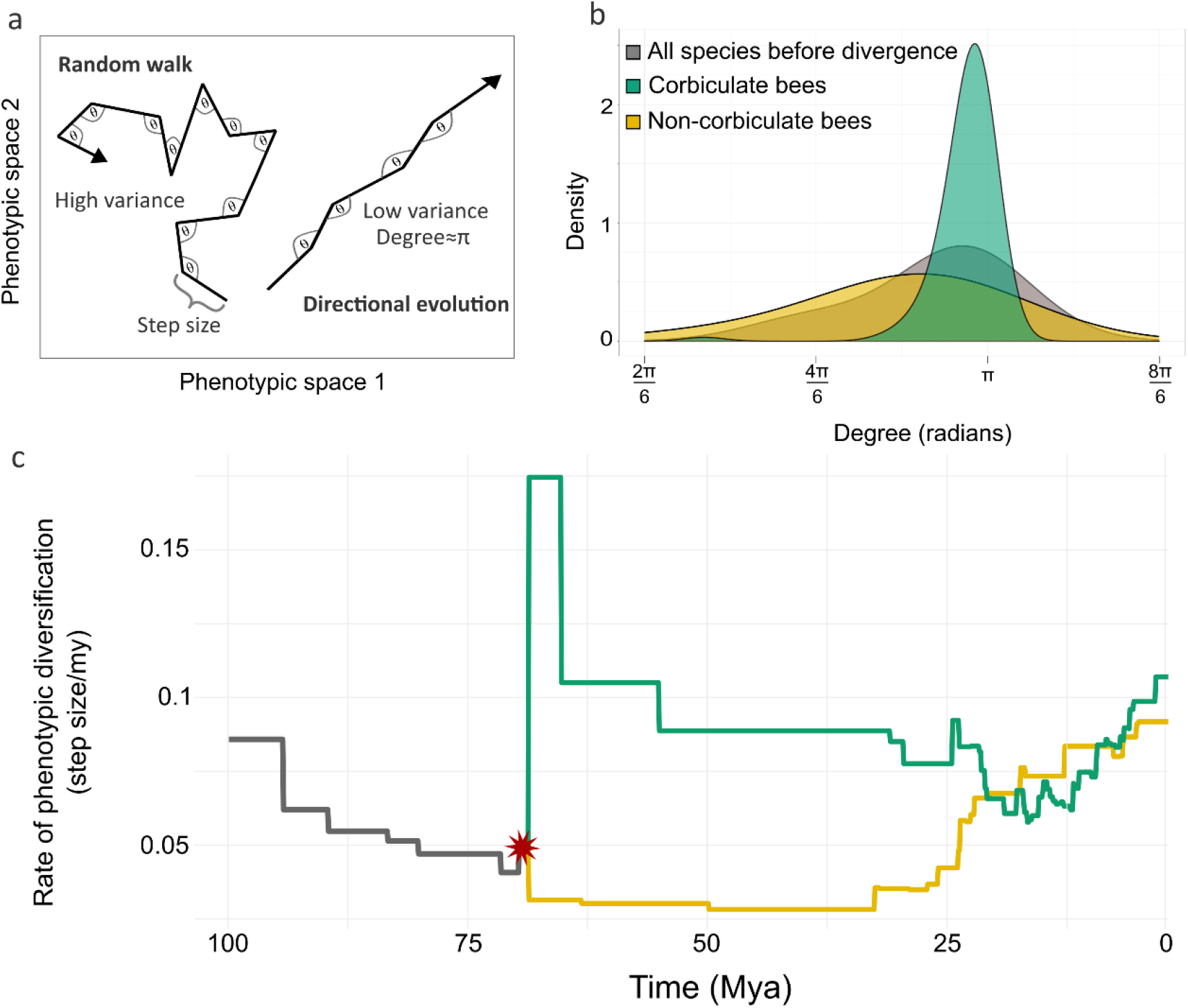
Analysis of assumed evolutionary trajectories in the phenotypic space of social complexity. Analyses are based on PCs 1-4. **(a)** Schematic representation of evolutionary routes under a directional evolution and a random walk. The mean and variance of the angles are used as measures for the directionality in the evolution of social complexity. The step sizes are used to measure the degree of phenotypic change along a branch. **(b)** Density plot of the turn degrees in the reconstructed evolutionary routes. **(c)** The phenotypic diversification rates across time before and after the inferred major evolutionary transition (indicated with a red asterisk). The gray line shows the average diversification of all species before the split; after the split, the rate of corbiculate bees is shown in green and that of the remaining taxonomic lineages is in yellow. The rate of phenotypic diversification is computed by dividing the lengths of the steps in the phenotypic spaces by the phylogenetic branch length.

Undergoing a major transition may remove biological constraints and open novel evolutionary opportunities, generating a phase in which the phenotypic space is rapidly explored and diversification is increased. Our findings are consistent with this notion: The rate of phenotypic diversification in the corbiculate bees substantially increased ∼70 mya, coinciding with the directed adaptive regime shift (green line in Fig. 5c). The phenotypic diversification of social complexity in the corbiculate bees quadrupled in rate for a short period, then decreased to about a half. In contrast, the remaining species show a rather constant low rate of diversification during this period. The later increase at ∼25 mya is attributed to the flexible and rapid changes in social complexity in these species occurring closer to the present, which is similar for corbiculate and non-corbiculate bees.

### Testing for selection signals in socially related genes

Lastly, we studied the selection patterns on several genes that have been repeatedly linked to the evolution of sociality in insects^35,66–69^, using the values of our main PCs as proxies for social complexity. We focused on insulin pathway-related genes (i.e., *IRS, InR, InR-2*), *TOR, syx1a*, and *Dunce* (Table S2). To detect signals of selection in these genes, we computed the molecular evolutionary rate (dN/dS values using the branch model of the ‘codeml’ program) for 10-18 species for which reliable genomic sequencing data was available (Table S3). We tested whether phylogeny, the social ladder categories^12,29^, or the main groups (A-D) in our dimensionality reduction analysis best explain the pattern of selection (Tables S4 and S5). For *InR-2*, we found a signal consistent with purifying selection in bumble bees (group C). For *IRS* and *InR* we detected a low signal for positive selection in *E. tridentata*. We found a similar low signal for *syx1a* in species from group A and for *TOR* in a single subsocial species *C. calcarata*. Lastly, we identified a strong signal of positive selection signal for *Dunce* in bumble bees (group C). Generally, our findings indicate that phylogeny and the phenotypic groups we identified better explain the selection patterns in focal social complexity genes than the social ladder classification (Tables S4 and S5).

These findings do not lend credence to the hypothesis that the selection on a few key social genes underpinned the evolution of social complexity across multiple independent origins of sociality in bees. Therefore, we next used linear regression analysis to test the correlation between PCs 1-4 and the dN/dS value of each gene, assessing whether the evolutionary rate of our focal genes is associated with quantitative indices of specific dimensions of sociality rather than with narrow classifications aimed at capturing the full social complexity phenotype. Remarkably, although we only tested six genes, we revealed significant correlation for two of them: *dunce* was correlated with PC4 (R^2^=0.34, *p*=0.01), and *syx1a* with PC1 (R^2^=0.32, *p*=0.02; Table S6).

## Discussion

We applied a bottom-up data-driven approach to analyze socially related traits, generating a phenotypic space of social complexity for 77 species of bees. By using multiple quantitative traits, we were able to avoid the semantic inconsistencies and *a priori* assumptions inherent in qualitative classifications, which currently dominate the research on the evolution of social complexity. The use of quantitative data also enabled rigorous analysis of social complexity and its evolution using methods that cannot be applied to qualitative data. With this novel approach, we identified clear clusters of species occupying defined areas in the phenotypic space of social complexity, corresponding to different social phenotypes (Figs. 2, S2-S5). Importantly, we identified substantial overlooked variability within each of these clusters. Our ancestral state reconstructions suggest a single major transition of increase in social complexity, which occurred in the common ancestor of honey bees, stingless bees, and bumble bees (Figs. 3-5). This transition into complex societies was followed by considerable phenotypic diversification (Fig. 5c). Non-corbiculate species with complex societies followed a different evolutionary trajectory than corbiculate bees. Notably, bumble bees, a group whose social complexity level is constantly debated^20,21,26,34^, align more closely with honey bees and stingless bees in our phenotypic space and are clearly separated from other species typically classified as primitively eusocial.

Amber fossils evidence from ∼45 mya is consistent with our findings suggesting that complex sociality in corbiculate bees dates back to their common ancestor^70,71^. These fossils from extinct corbiculate tribes show evidence of a clear worker caste, which had a barbed sting and reduced metasoma^70,71^. There are no other bee groups with fossil evidence of a morphological caste system. Additionally, no extant corbiculate bee species have reverted from social to solitary lifestyles (the *Psithyrus* Cuckoo bumble bees evolved a parasitic lifestyle that is not strictly solitary). This suggests that high levels of social complexity represent an evolutionary adaptive peak in the phenotypic landscape^72,73^, making a reversion to a less complex lifestyle unlikely^28^. Colonies of corbiculate bees have clear castes and the point-of-no-return is assumed to be crossed when unconditional differentiation of permanently unmated castes is established^13,21^. After crossing this threshold, a positive feedback process is assumed to support the further development of key social traits, such as finer division of labor, elaborated communication systems, and an increased colony size and survival^74–78^.

The corbiculate bees exhibit extensive phenotypic diversity in social complexity, which highlights the elaboration of trait interactions. Our results indicate that extant corbiculate diversity is the result of phenotypic diversification following an adaptive and directional evolutionary regime shift of approximately 70 mya (Figs. 4 and 5). According to conservative estimates from the fossil record, clade diversity among social corbiculate bees was higher in the past, experiencing at least a 50% reduction in social bee clades during the Eocene epoch (55-35 mya)^70,71^. This trend suggests that a major transition has opened novel evolutionary opportunities, driving the occupation of new niches in the social complexity phenotypic landscape. Such a macroevolutionary process likely involves overcoming structural, developmental, or ecological barriers, similar to the explosion of multicellular life following the development of sexual reproduction and division of labor^79,80^, or the rapid speciation observed after the colonization of remote islands^81,82^.

Sweat bees, orchid bees, and Xylocopinae bees exhibit lower levels of social complexity compared to corbiculate bees. We found evidence for repeated reversions to low levels of social complexity from group B to A, supporting earlier suggestions^23,26,83–88^. The reversion to lower social complexity suggests that these groups have not crossed the point-of-no-return. This notion is further supported by evidence for intra-species variability in social complexity with solitary and social lifestyles expressed under different environmental conditions^89–92^. The evolutionary increase in social complexity in these taxonomic groups was relatively continuous, without evidence for stepwise incremental changes. Two species, *E. tridentata* and *L. marginatum,* are typically used to describe independent origins of complex sociality (eusociality) in sweat bees and Xylocopinae. Our analyses confirm their relatively high social complexity but also raise questions about whether their social complexity and evolutionary trajectories are comparable to those of corbiculate bees. The social phenotype of these two species was inconsistent across our dimensionality reduction analyses (Figs. 2, S2, S4, S5), and their evolutionary trajectories in our phenotypic space differ from those of corbiculate bees, as well as from their close sister taxa. These findings likely reflect their unusual nesting biology; *E. tridentata* displays facultative social nesting with reproductive workers^93^, and *L. marginatum* lacks apparent queen-worker castes^94^. Additional investigations are needed to better describe their social complexity and robustly reconstruct their evolutionary history^36,95,96^.

The increased availability of life history and socially related data for a growing number of bee species enabled us to describe, for the first time, the phenotypic space of social complexity in bees and to study it quantitatively. Our data-driven approach circumvents the need to rely on explicit or implicit assumptions, facilitating unconstrained evolutionary inference^19,20,23,32–34,97^. Contrary to expectations of the social ladder perspective, our findings do not support a continuous^26^ or stepwise^12,29,98^ progression towards higher sociality across independent bee groups. Additionally, we did not find support for the notion that key social genes are selected across independent origins of sociality as would be expected in a narrow and restricted evolutionary trajectory towards sociality^29–31,99,100^. Our findings shed light on an important macroevolutionary process – traversal of a substantial phenotypic distance and subsequent diversification within corbiculate bees. Our framework can be expanded to additional animal groups, including primates and birds^6–8^, where research on the evolution of social complexity suffers from the same limitations as in social insects. Our approach is especially promising for studies on the molecular underpinnings of social complexity because it is more likely to capture the relationship between proxies of social complexity and genomic, epigenetic, or proteomic data, as demonstrated in the genomic analysis we conducted here.

## Methods

Unless stated otherwise, analyses described below were performed using R v.4.2.1^101^.

### Curating a dataset of social complexity traits in bees

We conducted an extensive literature survey using *Google Scholar* for any paper that includes relevant names of species and specific search words for each trait. Table S1 details the social traits that we used, their ecological context (e.g., field, lab, etc.), the keywords used to find relevant publications, and the references supporting the significance of the trait to social complexity. Overall, we surveyed >1000 articles, of which 219 were identified as providing relevant information, and were incorporated into our dataset. Based on the biology and life history of each focal species, we designed strict and consistent quality control guidelines to minimize the risk of including erroneous data that could introduce noise into our dataset (see Table S1 for details). For example, for average colony size to be included in the dataset, we required that it be measured for a healthy colony sampled during favorable conditions (e.g., in the spring) when food sources are abundant and the climate is suitable for colony growth. Similarly, colony longevity values were included only when measured in a natural setting or a field study, without intervention (e.g., supply of artificial food, disease control, etc.). In species with social polymorphism (i.e., when individuals can nest solitarily or socially), we used data for the most putatively advanced social form, unless stated otherwise.

We included in the dataset traits that are commonly used in the literature to describe various aspects of sociality, allowing us to acquire consistent data across multiple species. When determining which traits to include, we aimed to choose quantitative, informative, and comparable traits. For example, the percentage of workers with “active ovaries” can provide information on the degree of reproductive skew in a colony; however, the degree of ovary activation or development is often only qualitatively described and is measured differently across species; therefore, the latter trait was rejected. Furthermore, some traits are similar and complementary to one another, allowing us to obtain a comprehensive description of sociality aspects instead of assuming that one trait is indicative of another. For example, to measure the degree of reproductive skew in a colony, we used the percentage of mated workers, which indicates the totipotency of females, as well as the percentage of worker-produced males, which represents a more functional measure of the reproductive skew. Several traits that cannot be quantified but are important in the literature on insect sociality were coded as ordinal values (e.g., overlap of generations). Because our correlation matrix (Fig. 1c) showed that most traits are not highly correlated, the inclusion of similar traits does not inflate their importance in downstream analyses. To verify that no single trait biases our results, we conducted a sensitivity analysis in which we reanalyzed the data after removing every trait from the dataset (Fig. S11).

Our dataset includes species from all bee groups in which considerable social behavior has been reported (Apinae, Xylocopinae, Lasioglossum, Halictus, and Augochlorini). In choosing the species, we aimed to include as much variation as possible across the social complexity phenotype of bees. Therefore, our focus was more on sampling many different sociality phenotypes rather than extensively sampling the phylogeny. For example, stingless bees were found to be excellent candidates for diverse sampling and were indeed overrepresented in our dataset. On the other hand, many social species of sweat bees and Xylocopineae bees, which do not add additional information (i.e., they are similar to the species we included), were not overly sampled to avoid redundancy. Only two known species: *Hasinamelissa minuta* (Allodapine) and *Lasioglossum umbripenne* (Halictidae), might represent an informative sociality type^12,102^; however, insufficient data did not allow their inclusion. Furthermore, our sensitivity analyses with more equal sampling (Fig. S12) only reduced the variation within the phenotypic space but did not alter our results.

For each species in our dataset, we also provide the social complexity category using the common classification of Michener^12^. Because we wanted to minimize *a priori* assumptions, we did not include these classifications in our analyses but only used them as a reference to which we compared our findings. Overall, we gathered information on 77 species and 17 social traits, with 15% missing data in total. Four indices: colony average, colony maximum, colony foundation, and queen’s fecundity were log-transformed due to a high degree of variation that could skew the results (e.g., 1-80,000 for colony maximum). Additional details on the data curation procedure and the social traits and species included in our dataset are provided in Tables S1 and Dataset 1.

To impute missing data, we selected methods that offer sufficient statistical power while preserving the original data distribution. For example, we avoided approaches such as omitting all species with missing data or replacing missing data with the mean of the trait value, because this may bias or distort data distribution, respectively^103^. Based on our data structure, we used iterative PCA for imputation, which was found to perform well for data structures similar to ours, and specifically for PCA analyses^103^. For missing data imputation, we used the R package missMDA^104^. Each imputed data value was manually examined to verify that it was within the accepted ranges found in the literature on the relevant species. For example, we ensured that imputed values for colony longevity of known annual species are indeed of up to a year. Fourteen imputed entries (1% of the entries in our dataset) that did not meet this requirement or were not legitimate (e.g., negative values), were replaced manually with an approximated value based on the literature (e.g., species with reported colony longevity of “few months” without more accurate information was assigned a value of three months).

### Correlations and phylogenetic patterns of traits in the dataset

We used two methods to estimate the phylogenetic signal in the dataset: (i) Pagel’s λ^105^, which we estimated with maximum likelihood procedures using the phylosig function from the R package phytools^106^. Pagel’s λ value ranges from 0 (the trait evolved independently from phylogeny) to 1 (the trait fully depends on phylogeny and its evolution fits a neutral Brownian motion); (ii) Blomberg’s K^107^, which is not constrained to be between values 0 and 1, and K>1 implies that species are more similar to one another than expected under a Brownian motion process^107^. In other words, values higher than 1 indicate a constrained, lineage-specific trait. Additionally, the flexibility of Blomberg’s K allows us to capture the effects of changing evolutionary rates (e.g., non-Brownian motion processes)^108^. For both λ and K, we performed a likelihood-ratio test to evaluate the statistical significance of the phylogenetic signal^106^.

To test whether the different social traits are correlated, as implied by the social ladder model, we computed Spearman’s correlation coefficient between all trait pairs, while accounting for the shared evolutionary history between the species (i.e., non-independent data points). We first computed the phylogenetically independent contrasts (PICs) using the R package ape^109^, which implements the method described by Felsenstein^42^. This method computes the contrasts, or differences, in traits between each pair of sister taxa, generating a set of statistically independent values. We then computed Spearman’s correlation coefficients for these contrasts, allowing us to measure the strength and direction of the monotonic relationship between all pairs of variables. To account for the possibility of spurious correlations, we used the Benjamini–Hochberg method, which controls the false discovery rate (FDR) using sequential modified Bonferroni correction for multiple hypothesis testing^110^, as implemented in the p.adjust function. Only significant correlations (with a significance level of α=0.05) were considered as indicating correlated traits.

### Dimensionality reduction visualization

We used two procedures of dimension reduction, PCA and UMAP, to visualize and investigate the social phenotypic space of the species in our dataset. Principal component analysis (PCA) maximizes the variation in principal components and is useful for visualizing and quantifying a phenotypic space that captures the global structure of the dataset along a few major axes. In contrast, Uniform Manifold Approximation and Projection (UMAP) emphasizes local structures (e.g., within-group variation) in the dataset, while still accounting for the global structure (i.e., the between-group variation), and is more effective in visualizing tight clusters^111^. The combination of PCA and UMAP analyses provides us with a comprehensive reflection of a phenotypic space from our social complexity dataset (other dimensionality reduction techniques that focus primarily on local structures, such as t-SNE, would be less useful for this purpose).

For PCA, we used the R packages FactoMineR for visualization^112^, and MissMDA for performing PCA with missing data^104^. Before analysis, we standardized the trait values as z-scores. To correct the PCA for phylogeny, we applied pPCA using the phyl.pca function of the R package phytools^106^, which estimates the ancestral states of the traits using maximum likelihood under the λ model. This model allows variation in rates of evolution through time^105^ and was preferred over the passive Brownian motion model with equal rates. Finally, we used the broken stick method, as implemented in the R package PCDimension^63^, to determine the number of statistically significant PCs. This is done by randomly selecting *n* breakpoints (*n* being the number of variables in the dataset) from a uniform distribution; under this method, a PC is considered significant if it explains more variance than the randomly broken sticks. To understand the effect of traits on our phenotypic space, we examined the PCA loadings. Lastly, we computed the Euclidean space of the convex hull occupied by species in the phenotypic space of all four informative PCs and compared the area of corbiculate bees to the area of the remaining species. Analysis of UMAP was implemented in the R package umap^113^. Because phylogenetic correction has not yet been developed for UMAP, we established a phylogenetic procedure that uses the methods for PCA phylogenetic correction. This was achieved by taking the phylogenetically corrected PCA values from the first ten principal components and using them as input for the UMAP algorithm.

### Estimating the evolutionary trajectories of social complexity

Based on the common assumption that the ancestral lifestyle of bees was solitary^23,25^, we investigated the phenotypic trajectories by which social complexity increased throughout bee evolution. We performed ancestral state reconstruction using the ace function in the ape package^109^ and considered the PCA values (PCs 1-4) as describing the social complexity phenotype of the species, which together represent most of the variation of social complexity found in our dataset (∼70% of the variation). Because UMAP, unlike PCA, allows for data-dense regions to be stretched out in the representation, it is more challenging to interpret distances between species in the UMAP space, and therefore we did not use UMAP values for downstream analyses^114,115^. We set the root to the values of the solitary *Megachile rotundata*, which, unlike other solitary species in our dataset, belongs to a strictly solitary family and is more likely to represent a common phenotype of solitary bees. Fixing the root provides more statistical power because nodes deeper in the tree tend to be harder to estimate without *a priori* knowledge (e.g., fossils), and directional evolution, if exists, is difficult to capture^116^.

### Identifying shifts in social complexity

Maximum likelihood model fitting approaches have been widely used to quantify and compare evolutionary patterns in continuous traits (e.g., geometric morphometrics, functional morphology, biomechanical indices, etc.) based on categorical explanatory states (e.g., diet, habitat, life history, etc.)^117–121^. These methods are based on the adaptive evolution process of Ornstein–Uhlenbeck (OU)^73,122^. OU models are a modified Brownian motion process where the trait is attracted toward an optimum value. If the trait value at the root of the phylogeny is different from the optimum, the mean trait value of the lineages will tend to increase or decrease over time, eventually converging to around the optimum. In other words, OU models describe processes where trait values are constrained around one or several optima that can be considered adaptive peaks in a macroevolutionary landscape. However, this model-fitting approach still relies on *a priori* assumptions for species classification and, more importantly, forces qualitative transitions between states, which might not always be appropriate for reversible and small fluctuations in trait values. Recently developed methods^123–126^ have applied similar multi-peak OU models to detect shifts in the pattern of morphological evolution, creating an adaptive landscape of trait values that reflects Simpson’s framework of adaptive zones across multivariate trait spaces and multiple selective optima^127^. This approach is particularly relevant for our framework because it does not require an *a priori* hypothesis of shift locations (e.g., with changes in ecology or environments), nor classification of species and directionality of transitions between states (e.g., based on their social complexity level).

We used the PhylogeneticEM R package – a shift detection method that identifies phenotypic shifts along the phylogeny^126,128^. To account for trait correlations (i.e., interdependencies), PhylogeneticEM uses a simplified scalar OU (scOU) model which assumes that all traits evolve at the same rate towards their optimal values, if such optimum exists. This assumption facilitates the inference of the selection strength and drift rate for traits. The evolutionary trajectory of traits is modeled on a fixed, time-calibrated phylogenetic tree, with shifts in traits interpreted as responses to developmental, environmental, or phylogenetic changes. These shifts are integrated into the model as changes in the primary optimum. To determine the most supported number of shifts in the tree, the approach involves fixing the number of shifts and finding the best solution for that number, iterating this for various values of K (the number of shifts). An Expectation-Maximization (EM) algorithm^129^ is used for this process, which involves ancestral trait reconstruction and determining shift positions and magnitudes. The true number of shifts is estimated using a penalized likelihood criterion. We used two maximum likelihood methods to determine the number of shifts, both aim to balance between simple and weak models to complex overfitting models: LINselect evaluates how well each model predicts new data, instead of just how well it fits the data we have^130^. Djump (Jump Heuristic) looks for a jump in the model’s complexity that leads to a significant improvement in performance^131^.

### Estimation of directionality and phenotypic diversification in evolution

To explore the difference between species in the main sociality phenotypes (groups A-D; Fig. 2), we implemented a method from movement ecology to evaluate the directionality of the evolutionary routes of species in the phenotypic space. In the original method, the assumption is that movement is a correlated random walk, and movement directionality can be estimated by computing and analyzing the turning angle distributions of movement in a landscape^132^. Here, we used the high-dimensional phenotypic space of PCs 1-4 as the landscape and computed the turning angle between vectors along the evolutionary route of each species based on our ancestral reconstruction (Fig. S8). When the angle equals π (180**°**), we interpret this as no change in the direction of evolution; when the absolute value of the turning angle is higher or lower than π, evolutionary directionality is lower. We compared the distributions of angles between three groups of trajectories: all species before the divergence of corbiculate bees, corbiculate bees after divergence, and the remaining species after divergence. This comparison was to test whether putative trajectories that have crossed a major evolutionary transition display more directional evolution than trajectories that have not crossed it and show more flexible evolution. We performed an *F*-test to compare the variances between the groups and a *t*-test to compare the means.

We evaluated the phenotypic change in social complexity across time to estimate the rate of phenotypic diversification of extant taxonomic lineages. To compute phenotypic diversification, we again used the phenotypic space of PCs 1-4 and the evolutionary trajectories within this space. For each branch in the phylogeny, we calculated the Euclidean distance between the positions of the nodes defining the branch in the four-dimensional phenotypic space (i.e., the change in PC values; ΔPC). The distance between points represents the amount of phenotypic change along that branch. The rate of phenotypic diversification per million years is calculated by dividing the ΔPC by the temporal length of the branch, in mya:

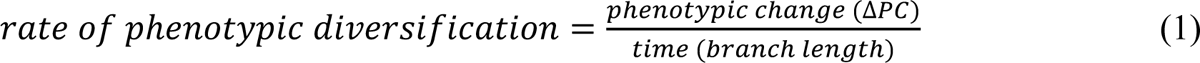

### Phylogenetic tree

We used species from across all the major social taxonomic lineages of bees: Apidae (honey bees–Apini, stingless bees–Meliponini, bumble bees–Bombini, orchid bees–Euglossini), Xylocopinae (Allodapini, Xylocopa and Ceratina), and Halictidae (sweat bees; including the tribe Augochlorini and the genus Lasioglossum and Halictus). Each of these groups is commonly considered to have evolved complex sociality (eusociality) independently^12,83^. Time-calibrated phylogenetic tree to be used in our analyses was taken from ‘beetreeoflife’^133^ and included all species in our dataset except *Scaptorigona postica*, which was replaced with its closest sister taxa *Scaptotrigona polysticta*. The phylogeny we used is based on the most comprehensive and up-to-date analysis of bee phylogenetics, constructed from a supermatrix of molecular data of more than 4500 species of bees across all families^51^.

## Acknowledgments

We thank Michael Schwarz for providing us with important data on *Exoneurella tridentata* and Igor Medici de Mattos for helping us translate papers from Portuguese.

## Code availability

All results and graphics are reproducible using the protocols detailed in the Methods. R code is deposited in GitHub and accessible at https://github.com/Greenbaum-Lab/bees.git

