## Supplementary material for "Data-driven analyses of social complexity in bees reveal phenotypic diversification following a major evolutionary transition": Dataset 1

| Species | CF | CM | CA | QL | CL | WS | QS | QW | PM | QF | OR | WM | SX | R | BC | NC | OG |
|---|---|---|---|---|---|---|---|---|---|---|---|---|---|---|---|---|---|
| *Apis mellifera* | 11800*[1]<br>C=126 | 80000[2] | 15000[3] | 2[4] | 5.6[5]<br>C=42 | 1.8[6] | 0.99[6] | 1[2] | 0.01[7]<br>C=57960<br>N=11<br>G | 1500[8,9] | 0.024*[10-12]<br>N>1700<br>C>7 | 0[2] | 0.002[9,13,14]<br>C=19 | 0.29[15]<br>C=5<br>N>400<br>G | 6[16] | 2[6] | 1[2] |
| *Apis cerana* | 10657[17]<br>C=20 | 40000[3] | 6900[18] | n.a. | n.a. | 1.77[6] | 1.08[6] | 1[2] | 0[7]<br>N=652<br>C=5<br>G | 960[6] | 0.112*[11,12]<br>N>150<br>C>5 | 0[2] | 0.001[17]<br>C=20 | 0.29[19]<br>C=4<br>G | 6[16] | 1[6] | 1[2] |
| *Apis dorsata* | 4000[20]<br>C=9 | 80000[3] | 36600[21] | n.a. | n.a. | 1.91[6] | 1.02[6] | 1[2] | 0[7]<br>N=660<br>C=4<br>G | 940[22]<br>C=20 | 0.188*[23]<br>N=1285<br>C=4 | 0[2] | 0.002[14]<br>C=15 | 0.29[24]<br>C=4<br>G | 6[16] | 1[6] | 1[2] |
| *Apis florea* | 2000[6] | 30000[6] | 6300[21] | n.a. | 1*[6] | 1.15[6] | 1.23[6] | 1[2] | 0[7]<br>N=564<br>C=4<br>G | 350[6] | 0.023[25,26]<br>N=47 | 0[2] | 0.01[14]<br>C=6 | 0.35[27]<br>C=5<br>G | 6[16] | 1[6] | 1[2] |
| *Ceratina australensis* | 1[28]<br>C>100 | 4[29] | 2[28]<br>C=36 | 1[28]<br>c>100 | n.a. | 4.8[28]<br>C≈70 | 1.02[28]<br>C≈70 | 0[28] | 0[29,30]<br>C=4<br>G | 1[29]<br>C=32 | 1[31] | 100[28] | 0.59[32]<br>C=150<br>N=405 | 0.79[29]<br>C=13<br>N=26<br>G | 1[28] | 0[28] | 1[28] |
| *Ceratina Calcarate** | 1[33]<br>C=563 | 1[33] | 1[33]<br>C=563 | 1[33] | 0[33] | 9.23[34,33]<br>N>500 | 1[33] | 0[33] | 100[33]<br>B | 1[33] | 1[31] | 100[2] | 0.44[35,34]<br>C=202<br>N=1182 | 0.48[36]<br>C=8<br>N≈75<br>G | 1[33] | 0[33] | 1[33] |
| *Exoneurella tridentata* | 1[37] | 37[37] | 10[37] | 4.5[37] | 15[37] | 4.98[37]<br>N=880 | 1.18[37]<br>N=62 | 1[37] | 1[37] | 1[37] | 1[37] | 8[37]<br>N=602 | 0.36[37]<br>N≈700 | 0.71[37]<br>C=27<br>N=318 | 5[37] | 0[37] | 1[37] |
| *Exoneura robusta* | 1[38]<br>C=40 | 8[38] | 2[38]<br>C=40 | 1[38]<br>C>100 | 0.5[39]<br>C=23 | n.a. | 1[40] | 0[38] | 29[38]<br>C=53<br>N>100<br>G | 1[38] | 1[41] | 100[38] | 0.83[42]<br>C=226<br>N=1857 | 0.39[38]<br>C=53<br>N>100<br>G | 5[43] | 0[43] | 1[38] |
| *Xylocopa pubescens* | 1[44]<br>C>100 | 2[44] | 2[44]<br>C>100 | 1[45]<br>C>100 | 0.16[44] | n.a. | 1[45] | 0[45] | 0[46]<br>C=20<br>B | 1[45] | 1[45] | 100[45] | 0.5*[47]<br>C=103<br>N≈480 | 0.48[48]<br>C=98<br>B | 1[45] | 0[44] | 1[44] |
| *Xylocopa virginica* | 1[49] | 5[49] | 2[49] | 2[49] | 8[50] | 3.16[51]<br>N=103 | 1.02[52] | 0[50] | 0[52]<br>B | 1[53] | 1[54] | 0[53] | 0.65[52,49]<br>C=38<br>N=393 | 0.15*[55]<br>N=118<br>G | 2[52] | 0[49] | 1[49] |
| *Ceratina cyanea* | 1[56] | 1[56] | 1[56] | 1[56] | 0[56] | n.a. | 1[56] | 0[56] | 100[57]<br>B | 1[56] | 1[56] | 100[56] | 0.38[56]<br>N=103 | 0.45[57]<br>C=2<br>G | 1[16] | 0[56] | 0[56] |
| *Exoneurella lawsoni* | 1[58] | 1[58] | 1[58] | 1[58] | 0[58] | n.a. | 1[58] | 0[58] | 100[58]<br>B | 1[58] | 1[58] | 100[58] | 0.58[58]<br>N=146 | n.a. | 4[58] | 0[58] | 0.5[58] |

| Species | CF | CM | CA | QL | CL | WS | QS | QW | PM | QF | OR | WM | SX | R | BC | NC | OG |
|---|---|---|---|---|---|---|---|---|---|---|---|---|---|---|---|---|---|
| *Braunsapis vitrea* | 1[59] | 3[59] | 2[59] | n.a. | n.a. | 5.52[59]<br>N=56 | 1.02[59]<br>N=98 | 0[59] | 0[59] | 1[59] | 1[59] | 0[59] | 0.76[59]<br>C=82<br>N=127 | n.a. | 4[59] | 0[59] | 1[59] |
| *Xylocopa violacea* | 1[60] | 1[60] | 1[60] | 0.41[60] | 0[60] | n.a. | 1[60] | 0[60] | 100[60]<br>B | 1[60] | 1[60] | 100[60] | 0.59[61]<br>N=229 | n.a. | 1[60] | 0[60] | 0[60] |
| *Euglossa viridissima* | 1[62] | 4[62] | 2[62] | 0.5[63] | n.a. | 2.05*[64]<br>N=10 | 1[62] | 0[62] | 30[62]<br>C=2<br>B | 1[62] | 1[65] | 100[62] | 0.6[64]<br>C=27<br>N=360 | 0.52*[66] | 3[62] | 0[62] | 1[62] |
| *Euglossa townsendi* | 1[67] | 6[67] | 2[67] | 0.49[67] | 0.18[67]<br>C=5 | n.a. | 1[67] | 0[67] | 0[67]<br>C=5<br>B | 1[67] | 1[67] | 100[67]<br>N=6 | 0.6[67]<br>N=58 | n.a. | 3[67] | 0[67] | 1[67] |
| *Eulaema meriana* | 1[68] | 2[68] | 2[68] | 0.33[63] | 0.05[68]<br>C=1 | 2.23*[68]<br>N=46 | 1[68] | 0[68] | 100[68]<br>C=1<br>B | 1[68] | 1[65] | 100[68] | n.a. | n.a. | 1[68] | 0[68] | 0[68] |
| *Euglossa cordata* | 1[69] | 6[69] | 2[69] | 0.52[69] | 0.08[69]<br>c=11 | 4.47[69]<br>N=54 | 1.01[69]<br>N=20 | 0[69] | 0[70]<br>C=13<br>G | 1[69] | 1[70] | 100[69] | n.a. | 0.56[70]<br>C=13<br>G | 3[69] | 0[69] | 1[69] |
| *Eulaema nigrita* | 1[71] | 6[71] | 4[71] | 0.21[71,72] | 0.11[71]<br>C=9 | 4.53*[71]<br>N=8 | 1[71] | 0[71] | 100[71]<br>C=9<br>B | 1[71] | 1[71] | 100[71] | n.a. | 0.75*[71]<br>B | 1[71] | 0[71] | 0[71] |
| *Euglossa melanotricha* | 1[65] | 3[65] | 3[65] | 0.22[73]<br>C=30 | 0.1[65]<br>C=26 | n.a. | 1[73] | 0[73] | 30[65,73]<br>C=56<br>B | 1[65] | 1[65] | 100[65] | 0.64[65,74]<br>C=48<br>N=504 | 0.66[74]<br>C=18<br>G | 3[65] | 0[65] | 1[65] |
| *Euglossa atroveneta* | 1[75] | 10[75] | 6[75] | 0.58*[75] | 0.33[75]<br>C=9 | n.a. | 1[75] | 0[75] | n.a. | 1[75]<br>C=9 | 1[75]<br>C=9 | 100[75]<br>C=9 | n.a. | 0.62*[75] | n.a. | 0[75] | 1[75] |
| *Euglossa dodsoni* | 1[76] | 1[76] | 1[76] | n.a. | 0[76] | n.a. | 1[76] | 0[76] | 100[76]<br>B | 1[76]<br>C=8 | 1[76]<br>C=8 | 100[76]<br>C=8 | 0.53[76]<br>C=8 | n.a. | 1[76] | 0[76] | 0[76] |
| *Euglossa hyacinthina* | 1[77] | 5[77] | 2[77] | 0.46[78] | n.a. | 2.28[77,79]<br>N=80 | 1[77] | 0[77] | 100[77]<br>C=11<br>B | 1[77]<br>C=11 | 1[77]<br>C=11 | 100[77]<br>C=11 | 0.69[77,79]<br>C=55<br>N=190 | n.a. | 1[77] | 1[80] | 1[77] |
| *Bombus terrestris* | 1[81] | 400[81] | 200[82] | 1[81] | 0.33[83] | 11.3*[84]<br>C>40<br>N>5000 | 1.76[85]<br>C=28<br>N=868 | 0[16] | 3.4[7]<br>C=47<br>N=3749<br>G | 17[86]<br>C=18 | 1[16] | 0[16] | 0.23[86]<br>C=18<br>N=3178 | 0.75[7]<br>N=45<br>G | 5[16] | 1.5[88] | 1[88] |
| *Bombus Impatiens* | 1[81] | 756[89] | 450[16] | 1[81] | 0.19[90] | 9.96[91]<br>N=45 | 1.5[90]<br>C=11<br>N=239 | 0[16] | 0[92]<br>C=18<br>N=324<br>B | 10[90]<br>C=11 | 1[93] | 0[16] | 0.38[90,94]<br>C>15<br>N>2000 | 0.75[7]<br>C=10<br>N=180 | 5[16] | 1[16] | 1[88] |

| Species | CF | CM | CA | QL | CL | WS | QS | QW | PM | QF | OR | WM | SX | R | BC | NC | OG |
|---|---|---|---|---|---|---|---|---|---|---|---|---|---|---|---|---|---|
| *Bombus atratus* | 50[88] | 1605[95] | 480[96,97] | 1[98] | 5[98,88] | 7.7[84]<br>N=30 | 1.29[99]<br>N=9 | 0[16] | 90[88]<br>B | 16[97,100]<br>C=8 | 1[96] | 1[97] | 0.3[98,96]<br>C=32 | n.a. | 2[97] | 1.5[97] | 1[88] |
| *Bombus hortorum* | 1[81] | 128[81] | 100[81] | 1[81] | 0.17[83] | 9.1[84]<br>N=158 | 1.17[101]<br>N=10 | 0[16] | n.a. | 20[16] | 1[31] | 0[16] | n.a. | 0.75[102]<br>B,S | 2[103] | 1.5[88] | 1[88] |
| *Bombus polaris* | 1[81] | 21[104] | 16[105] | 1[81] | 0.16[105] | n.a. | 1.24[101]<br>N=8 | 0[16] | 0[105]<br>C=8<br>B | 15[104] | 1[31] | 0[16] | 0.35[105]<br>C=8<br>n>50 | n.a. | 5[105] | 1[88] | 1[88] |
| *Bombus morio* | 1[81] | 525[106] | 300[106] | 1[81] | 0.38[107] | 9.73[107]<br>C=2<br>N=135 | 1.3[101]<br>N=10 | 0[16] | n.a. | n.a. | 1[31] | 0[16] | 0.34[107]<br>C=2<br>N=71 | 0.75[88]<br>B,S | 2[108] | 1.5[88] | 1[88] |
| *Bombus pratorum* | 1[81] | 130 | 100[109] | 1[81] | 0.21[81] | 7.2[84]<br>N=189 | 1.22[101]<br>N=10 | 0[16] | n.a. | 7[88,110] | 1[31] | 0[16] | n.a. | 0.75[102]<br>N=35<br>G | 5[103] | 1[88] | 1[88] |
| *Bombus lapidarius* | 1[81] | 400[88] | 300[88] | 1[81] | 0.3[111] | 7.9[84]<br>N=385 | 1.45[101]<br>N=10 | 0[16] | n.a. | n.a. | 1[31] | 0[16] | n.a. | 0.75[102]<br>N=20<br>G | 5[103] | 2[88] | 1[88] |
| *Bombus lucorum* | 1[81] | 400[88] | 130[16,112] | 1[81] | 0.25[111] | 7.05[84]<br>N=269 | 1.11[101]<br>N=9 | 0[16] | n.a. | 3*[113]<br>C=12 | 1[31] | 0[16] | 0.37[112]<br>C=36 | 0.75[102]<br>N=20<br>G | 5[103] | 1.5[88] | 1[88] |
| *Bombus muscorum* | 1[81] | n.a. | 50[16] | 1[81] | n.a. | 11.05[84]<br>N=507 | n.a. | 0[16] | n.a. | n.a. | 1[31] | 0[16] | n.a. | n.a. | 2[103] | 1[88] | 1[88] |
| *Bombus Pascuorum** | 1[81] | 180[109] | 60[16] | 1[81] | 0.42[111] | 9.1[84]<br>N=255 | n.a. | 0[16] | n.a. | 9[16] | 1[31] | 0[16] | 0.61[114]<br>C=7<br>N=217 | 0.75[102]<br>B,S | 2[103] | 1[88] | 1[88] |
| *Bombus hypnorum* | 1[81] | 150[115] | 50[115] | 1[81] | 0.22[116] | 5.51*[117] | 1.44[101]<br>N=10 | 0[16] | 15.4[7]<br>C=28<br>N=1426 | 2*[116]<br>C=9 | 1[31] | 0[16] | 0.34[115]<br>C=24 | 0.69[7]<br>G | 5[118] | 1[88] | 1[88] |
| *Bombus ignitus* | 1[81] | 240[119] | 107[119] | 1[81] | 0.35[120] | 11.64<br>N=89 | 1.21[101] | 0[16] | 4.1[121]<br>C=5 | 5[120]<br>C=12 | 1[31] | 0[16] | 0.1*[122,123]<br>C≈44 | 0.75[124]<br>S | 5[125] | 1.5[126] | 1[88] |
| *Bombus fervidus* | 1[81] | 300[88] | 250[88] | 1[81] | n.a. | n.a. | 1.17[101]<br>N=9 | 0[16] | n.a. | 8[110]<br>C=7 | 1[31] | 0[16] | n.a. | n.a. | 5[110] | 1[88] | 1[88] |
| *Bombus hypocrita* | 1[81] | 260[127] | 45[120] | 1[81] | 0.31[120] | 9.75[123]<br>N=138 | 1.35[101] | 0[16] | 21[126]<br>C=1<br>B | 4[120] | 1[31] | 0[16] | 0.185[126]<br>C=2 | n.a. | 5[126] | 2[126] | 1[88] |
| *Bombus occidentalis* | 1[81] | n.a. | n.a. | 1[81] | n.a. | n.a. | 1.38[101]<br>N=10 | 0[16] | n.a. | 8[88] | 1[31] | 0[16] | n.a. | n.a. | 5[125] | 1[88] | 1[88] |
| *Bombus melanopygus* | 1[81] | 133 | 50 | 1[81] | 0.33*[128] | n.a. | 1.36[101]<br>N=10 | 0[16] | 19.1*[7]<br>C=15<br>N=112 | n.a. | 1[31] | 0[16] | 0.12[94]<br>C=17<br>N=1700 | 0.75[7] | 5 | 1[88] | 1[88] |

| Species | CF | CM | CA | QL | CL | WS | QS | QW | PM | QF | OR | WM | SX | R | BC | NC | OG |
|---|---|---|---|---|---|---|---|---|---|---|---|---|---|---|---|---|---|
| *Tetragonisca angustula* | 500[129] | 8000[129] | 5000[129] | 5[130] | 8.15[131]<br>C=141 | 3.2[84] | 1.85[132] | 1[16] | 0[133]<br>C=3 | 154[134]<br>C=4 | 1[135] | 0[16] | 0.01[136]<br>C=3 | n.a. | 3[129] | 5[a] | 1[129] |
| *Melipona quadrifasciata* | 300[129] | 1500[129] | 900[129] | 3[137] | n.a. | 4.13[84]<br>N=60 | 0.91[132] | 1[16] | 64[7]<br>N=47<br>C=2<br>G | 24[138]<br>C=1 | 1[135] | 0[16] | 0.47[129] | 0.83[139]<br>C=3<br>N=39<br>G | 3[129] | 5[a] | 1[129] |
| *Frieseomelitta varia* | n.a. | 1600[129] | 1200[129] | n.a. | n.a. | 1.56[84] | 1.85*[132] | 1[16] | 0[129]<br>P | n.a. | 0[135] | 0[16] | n.a. | n.a. | 3[129] | 2[a] | 1[129] |
| *scaptotrigona postica* | 2000[129] | 50000[129] | 15000[129] | n.a. | n.a. | 2.24[84]<br>N=52 | 1.25*[132] | 1[16] | 14.7[7]<br>N=284<br>C>3<br>G | 543[136,140]<br>C=5 | 1[135] | 0[16] | n.a. | 0.7[141,142]<br>C=11<br>N=204<br>G | 3[129] | 5[a] | 1[129] |
| *Melipona beecheii* | 300[129] | 3000[129] | 800[129] | 3[129] | 15[131]<br>C=11 | 1.92[143]<br>N=80 | 0.85[132] | 1[16] | 0[7]<br>N=108<br>C=13<br>G | 18*[144]<br>C=1 | 1[145] | 0[16] | 0.37[146]<br>C=6 | 0.68[141]<br>C=10<br>N=99<br>G | 3[129] | 5[a] | 1[129] |
| *Austroplebeia australis* | 1000[129] | 5000[129] | 3000[129] | n.a. | n.a. | 2.81[147]<br>n>100 | 1.38[132] | 1[16] | 7[7]<br>N=94<br>C=1<br>G | 39[148]<br>C=1 | n.a. | 0[16] | n.a. | 0.78[149]<br>G | 3[129] | 5[a] | 1[129] |
| *Apotrigona nebulata* | 195[129] | 2000[129] | 1700[129] | n.a. | 5.5[129] | n.a. | n.a. | 1[16] | n.a. | n.a. | n.a. | 0[16] | 0.001*[129] | n.a. | 3[129] | 5[a] | 1[129] |
| *Cephalotrigona capitata* | n.a. | 1000[129] | 700[129] | n.a. | 23[131]<br>C=17 | n.a. | 1.3[132] | 1[16] | n.a. | n.a. | n.a. | 0[16] | n.a. | n.a. | 3[129] | 5[a] | 1[129] |
| *Friesella schrottkyi* | 300[129] | 2500[129] | 1400[129] | n.a. | n.a. | 2.39[84] | n.a. | 1[16] | 0[150]<br>C=14<br>P | 53[151]<br>C>2 | n.a. | 0[16] | n.a. | n.a. | 3[129] | 4.5[a] | 1[129] |
| *Geotrigona mombuca* | n.a. | 3000[129] | 2500[129] | n.a. | n.a. | 1.42[84] | n.a. | 1[16] | 0[137]<br>C=3<br>B | n.a. | n.a. | 0[16] | n.a. | n.a. | 3[129] | 5[a] | 1[129] |
| *Hypotrigona gribodoi* | 100[129] | 750[129] | 450[129] | 4127 | 5.5[129] | n.a. | n.a. | 1[16] | n.a. | 50[129] | n.a. | 0[16] | n.a. | n.a. | 3[129] | 4[a] | 1[129] |
| *Lestrimelitta limao* | 200*[153] | 7000[129] | 4500[129] | n.a. | 10[154]<br>C=4 | 1.44[84] | 1.38[132] | 1[16] | n.a. | 670[129] | 0.32[155] | 0[16] | n.a. | n.a. | 3[129] | 5[a] | 1[129] |
| *Melipona fasciata* | 200[129] | 2500[129] | 1000[129] | n.a. | 10 | 1.5[84] | 0.9*[132] | 1[16] | 1[137]<br>B | n.a. | 1[156] | 0[16] | 0.5*[129] | n.a. | 3[129] | 5[a] | 1[129] |
| *Melipona fasciculata* | 300[129] | 3000[129] | 600[129] | n.a. | n.a. | 1.13[84] | 0.85*[132] | 1[16] | 0[157]<br>C=9<br>B | n.a. | 1[156] | 0[16] | n.a. | 0.75*[158] | 3[129] | 5[a] | 1[129] |
| *Melipona favosa* | 60[129] | 700[129] | 400[129] | 3[129] | n.a. | n.a. | 0.75[132] | 1[16] | 94.5[7]<br>N=604<br>C=4<br>G | 10.1[159]<br>C=6 | 1[156] | 0[16] | 0.26[129]<br>C>78 | n.a. | 3[129] | 5[a] | 1[129] |

| Species | CF | CM | CA | QL | CL | WS | QS | QW | PM | QF | OR | WM | SX | R | BC | NC | OG |
|---|---|---|---|---|---|---|---|---|---|---|---|---|---|---|---|---|---|
| *Melipona flavolineata* | n.a. | 3300[129] | 1500[129] | n.a. | n.a. | 2.02[84] | 0.87*[132] | 1[16] | n.a. | n.a. | 1[156] | 0[16] | n.a. | 0.75*[158] | 3[129] | 5[a] | 1[129] |
| *Melipona fuliginosa* | 250[129] | 600[129] | 383[129] | n.a. | n.a. | n.a. | 0.9[132] | 1[16] | n.a. | n.a. | 1[156] | 0[16] | n.a. | n.a. | 3[129] | 5[a] | 1[129] |
| *Melipona bicolor* | 150[129] | 800[129] | 425[129] | 7[129] | n.a. | n.a. | 0.84[132] | 1[16] | 54*[160]<br>C=1<br>B | 30[161]<br>C=1 | 1[156] | 0[16] | 0.5[162]<br>C=18 | 0.64*[129]<br>C=14<br>G | 3[129] | 5[a] | 1[129] |
| *Melipona marginata* | 160[129] | 2500[129] | 1330[129] | n.a. | n.a. | n.a. | 0.81[132] | 1[16] | 37.1[7]<br>N=41<br>C=3<br>G | 28[163]<br>C=2 | 1[156] | 0[16] | 0.6[129] | 0.7[139]<br>C=5<br>N=51<br>G | 3[129] | 5[a] | 1[129] |
| *Melipona scutellaris* | n.a. | 2000[129] | 1500[129] | n.a. | 15[154]<br>C=6 | 1.78[84] | 0.9[132] | 1[16] | 23[164]<br>C=45<br>G | 21.3[165]<br>C=1 | 1[156] | 0[16] | n.a. | 0.77[139]<br>C=9<br>N=89<br>G | 3[129] | 5[a] | 1[129] |
| *Nannotrigona testaceicornis* | 200[129] | 3000[129] | 2500[129] | n.a. | 10[154]<br>C=7 | 1.6[84] | 1.11*[132] | 1[16] | n.a. | 16*[166]<br>C=1 | 0.44[167] | 0[16] | 0.005[129]<br>C>2 | n.a. | 3[129] | 4[a] | 1[129] |
| *Plebeia remota* | 800[129] | 5000[129] | 2900[129] | 3[129] | 8.15*[168] | n.a. | n.a. | 1[16] | 3[137]<br>C=5<br>G | 220[169] | 0.66[135] | 0[16] | 0.003[170,162]<br>C=8 | 0.79[139]<br>C=7<br>N=79<br>G | 3[129] | 5[a] | 1[129] |
| *Plebeia droryana* | 200[129] | 10000[129] | 5000[129] | n.a. | 10[154]<br>C=11 | 2.21[84] | 0.97*[132] | 1[16] | 15.2[7]<br>N=300<br>C=15<br>G | 80[129] | n.a. | 0[16] | n.a. | n.a. | 3[129] | 5[a] | 1[129] |
| *scaptotrigona pectoralis* | 1400[129] | 5200[129] | 3600[129] | n.a. | 23[131]<br>C=33 | n.a. | 1.14[132] | 1[16] | n.a. | n.a. | n.a. | 0[16] | n.a. | 0.71[171]<br>C=7<br>G | 3[129] | 4[a] | 1[129] |
| *Schwarziana quadripunctata* | 500[129] | 2500[129] | 1500[129] | n.a. | n.a. | n.a. | 1.17[132] | 1[16] | 0[7]<br>N=314<br>C=16<br>G | n.a. | 0.61[162] | 0[16] | 0.03[162]<br>C=20 | 0.79[142]<br>C=4<br>G | 3[129] | 5[a] | 1[129] |
| *Tetragonula carbonaria* | 2500[129] | 11000[129] | 6750[129] | n.a. | n.a. | n.a. | n.a. | 1[16] | 0[7]<br>N=20<br>C=1<br>G | 300[172]<br>C=7 | 0[172] | 0[16] | n.a. | 0.75[173]<br>C=5<br>N=48<br>G | 3[129] | 5[a] | 1[129] |

| Species | CF | CM | CA | QL | CL | WS | QS | QW | PM | QF | OR | WM | SX | R | BC | NC | OG |
|---|---|---|---|---|---|---|---|---|---|---|---|---|---|---|---|---|---|
| *Trigona spinipes* | 5000[174] | 53000[174] | 39000[174] | n.a. | 10[154]<br>C=2 | 2.7*160<br>N=20 | 1.29*[132] | 1[16] | n.a. | n.a. | 0.33[176] | 0[16] | n.a. | 0.75[177]<br>C=43<br>G | 3[129] | 5[a] | 1[129] |
| *Trigona fulviventris* | 2000[129] | 10000[129] | 5750[129] | n.a. | 23[131]<br>C=10 | n.a. | 1.31[132] | 1[16] | n.a. | n.a. | n.a. | 0[16] | n.a. | 0.68[142]<br>C=7<br>G | 3[129] | 5[a] | 1[129] |
| *Megalopta genalis* | 1[178] | 11 [178] | 3[178] | 0.65*[179] | 0.11[180]<br>C=170 | 9.11[181]<br>N=61 | 1.08[182]<br>N=46 | 0[178] | 3[183]<br>C=30<br>G | 1[178] | 1[178] | 75[178]<br>C=36 | 0.46[184]<br>C=970<br>N=2500 | 0.75[183]<br>C=30<br>G | 2[178] | 0[178] | 1[178] |
| *Lasioglossum baleicum* | 1[185] | 7[185] | 2[185] | 1[185] | 0.12[186]<br>C≈35 | 1[186]<br>N=76 | 1.05[185]<br>C=22 | 0[185] | 0*[187]<br>C≈30 | 1[185] | 1[185] | 57[185]<br>C=50 | 0.8[185,186]<br>C=85 | 0.7[187]<br>C=33<br>G | 2[185] | 0[185] | 1[185] |
| *Lasioglossum marginatum* | 1[16] | 486*[16] | 280[16] | 6[16] | 6[16] | n.a. | 1[188] | 0[16] | n.a. | n.a. | 1[16] | 0*[16] | 0.3[188]<br>C=9 | n.a. | 2[16] | 2[16] | 1[16] |
| *Halictus Ligatus** | 1*[189] | 20[189] | 4[189] | 1[189] | 0.2[189] | 6.1[190]<br>N=378 | 1.14[189]<br>N=216 | 0[189] | 50*[191]<br>C=27<br>N=106<br>G | 1[189] | 1[189] | 51[189]<br>N=264 | 0.65[192]<br>C=58<br>N=217 | 0.42[191]<br>C=27<br>N=106<br>G | 2[189] | 1[189] | 1[189] |
| *Lasioglossum villosulus* | 1[193] | 1[193] | 1[193] | 0.75[193] | 0[193] | 3.65*[193]<br>N=402 | 1.07*[193]<br>N=402 | 0[193] | 100[193] | 1[193] | 1[193] | 100[193] | 0.66*[193]<br>N=213 | n.a. | 1[193] | 0[193] | 1[193] |
| *Lasioglossum zephyrum* | 1[16] | 45[16] | 14[194] | 1[194] | 0.29[195] | 0.5*[196]<br>N=105 | 1.06[194,196]<br>N>156 | 0[16] | 15[197]<br>C=14<br>N=176<br>G | 1[16] | 1[16] | 8[16] | 0.73*[195]<br>C=27<br>N=669 | 0.7*[198]<br>C=20<br>G | 2[16] | 1[16] | 1[16] |
| *Habropoda laboriosa* | 1[199] | 1[199] | 1[199] | 0.08*[200] | 0[199] | 2.46[201]<br>N=48 | 1[199] | 0[199] | 100[199] | 1[199] | 1[199] | 100[199] | n.a. | 0.75*[202]<br>B | 1[199] | 0[199] | 0[199] |
| *Dufourea novaeangliae* | 1[203] | 1[203] | 1[203] | 0.08[203] | 0[203] | n.a. | 1[203] | 0[203] | 100[203] | 1[203] | 1[203] | 100[203] | n.a. | n.a. | 1[203] | 0[203] | 0[203] |
| *Megachile rotundata* | 1[204] | 1[204] | 1[204] | 0.16[204] | 0[204] | 2.4*[205]<br>N=500 | 1[204] | 0[204] | 100[204] | 2[204] | 1[204] | 100[204] | 0.35[205]<br>N~1500 | 0.65[206]<br>N=71<br>G | 1[204] | 0[204] | 0[204] |

C=colonies

N=individuals

G- data obtained via genetic test (allozymes, microsatellites, DNA fingerprints).

B- data obtained via behavioral observations.

S- data obtained via spermatheca count, which was suggested as a good estimator[102].

a- data is from [155,207–210]

***Apis mellifera:*** Data were taken without a clear distinction between the numerous subspecies, excluding *Apis mellifera capensis*. Whenever there was data for more than one subspecies, we calculated the average; CF-number of swarms and not colonies.
***Apis cerana*:** Taken from bees 1-10 days old.
***Apis dorsata:*** Data on worker ovarioles were taken from queenless colonies. However, the percentage of workers with slightly active ovaries and visible ovarioles was low in these colonies (14%, and only four individuals with highly active ovaries) - similar to the situation in queenright colonies39.⋰.
***Apis florea:*** All colony members leave the nest in multiple swarming events after a year.
***Ceratina calcarate:*** We refer to subsocial colonies.
***Xylocopa pubescens:*** Personal comment in [47].
***Xylocopa virginica:*** The study was conducted during the nestmate provisioning phase, prior to brood provisioning, so it is possible that the social unit that was tested was not entirely complete.
***Eulaema nigrita:*** WS-measured for metasoma width; R-behavioral observations suggest that nestmates stay in their nest and are of the same generation. Because Euglossini are monandrous, we assumed a 0.75 relatedness between full sisters[212].
***Euglossa atroveneta:*** QL-minimal queen longevity was inferred from a coexistence of mother and daughter in reactivated nest (colony longevity) of four months added to two month of brood development that the mother had to wait for brood to emerge and another month of provisioning; R-no specific data but social nests comprised of mother-daughter (r=0.5) or daughter-daughter (r=0.75). We assumed monandry as suggested for Euglossini [212] and took an average.
***Euglossa viridissima:*** WS-Based on body weight after emergence; R-based on 4 matrifilial colonies (3 mother-daughter, 1 mother-2 daughters) and a single mating.
***Eulaema meriana*:** Assumption based on brood cell volume of 6 nests: the uniformity in size of brood cells may account for the fact that body size variation in *E. meriana* appears to be negligible, at least locally[68].
***Bombus atratus:*** Species name was recently changed *to Bombus paulensis*; CF-colony reproduction by a lone foundress or swarming of several gynes with several workers from other colonies. We estimated the average (1-80); CL-colony can live for a few months like other Bombus spp. and up to 10 years. In addition, gynes can go back to their natal nest and supersede the queen, thus maintaining a perennial colony. We took an average (0.5-10).
***Bombus locorum:*** FQ was calculated by 32.6 (workers) + 121.2 (males) in approximately 60 days = 2.7.
***Bombus hypnorum*:** FQ-was calculated by number of males and workers produced by queen (153) divided by her reproductively active time (≈77 days); WS-data were given as plot and categorized workers to three behavioral groups. We extracted the sample size, mean and sd for each group and simulated the size for all workers combined assuming normal distribution, then calculated the mean and s.d. by ten iterations of random sampling that averaged 5.51 (4.51-6.21).
***Bombus ignitus:*** Colonies were reared in artificial conditions with different temperature\humidity but with consistent results.
***Bombos pascourom*:** Synonym to *bombus agrorum.*
***Exoneurella tridentata:*** data for WS and SR was taken from plot. WS was calculated for body size (not head\thorax weight).
***Bombus melanopygus:*** CL-two generations (8 months in total) divided by 2; PM-estimation of PM is based on the screening of 1,125 males using the Mendelian inherited abdominal pile-coloration marker. Since the power to detect workers' sons differed among colonies, PM was estimated using a maximum likelihood approach [7].
***Lestrimelitta limao:*** Estimation for incipient stage of swarming.
***Plebeia remota*:** Survival rate similar to *tetragonisca angostula* but no exact data.
***Trigona spinipes:*** Data taken from foraging bees.
***Geotrigona mombuca*:** Estimated at 1.65.
***Scaptotrigona postica:*** Data from sister taxa *Scaptotrigona Mexicana.*
***Nannotrigona testaceicornis:*** Data from sister taxa *Nannotrigona perilampoides.*
***Trigona spinipes:*** Data from sister taxa *Trigona corvina.*
***Frieseomelitta varia:*** Data from sister taxa *Frieseomelitta nigra.*
***Plebeia droryana:*** Data from sister taxa *Plebeia frontalis.*
***Melipona fasciata:*** QS - data from sister taxa Melipona scutellaris; SX-uncertain identity[129].
***Melipona fasciculata:*** QS-data from sister taxa *Melipona beechelii;* R-estimated from single mated queen.
***Melipona flavolineata:*** Data from sister taxa *Melipona rufiventris;* R-estimated from single mated queen.
***Melipona bicolor:*** Data was estimated to be between 27-82, and we took the median 54.
***Melipona beecheii:*** Observations were conducted for a week.
***Nannotrigona testaceicornis:*** Observations were done for 9 days.

***Melipona scutellaris:*** Regular sized queen has 12 ovarioles and a miniature queen has 4-8. We took an average.
***Apotrigona nebulata:*** Gruter [129] gives a value of ~0% queens and 10.5% males with no further available data. We assume from this a very low number of queens produced and use a more plausible value of 0.02% queens (similar to honeybees). Thus sex ratio is 0.02/(10.5+0.02) =0.001
***Melipona bicolor:*** Can be polygynous (0.75) and monandrous (0.53). We took an average.
***Megalopta genalis:*** QL-in observation nests (represents maximum longevity).
***Lasioglossum baleicum:*** Inferred from dissection of sterile workers.
***Lasioglossum marginatum***: CM- there are suggestions to higher numbers up to 897, but with uncertainty if it is only workers or combined with gynes+males. Thus, we took the conservative value of putative workers only; WM-in natural conditions it seems that workers do not mate and have undeveloped ovaries. However, on certain conditions females can mate and become queens [188].
***Halictus ligatus*:** A species with high degree of intraspecific polymorphism. We used data of northern populations (see references in table); CF-small proportion of nests have 2-3 foundresses; R-in this species workers produces mostly gynes rather than males, thus we give the value for gynes produced by workers.
***Lasioglossum villosulus***: WS-based on the average of two separate generations; QS-size differences of two generations, but no reproductive skew between them; SX-taken from laboratory experiment but confirms data from the field.
***Lasioglossum zephyrum:*** WS-lab conditions; SX-lab conditions; R-the limits from the sample used of 20 nests are 0.64<0.82<1. While these limits are consistent with the true value being 0.75 (that expected under male-haploidy if each nest results from the reproduction of a single, once-mated female), the occurrence of some nests with three or more genotypes shows that nest makeup is more complex than this, so that a slightly lower value, 0.7, is more plausible.
**Habropoda laboriosa:** Data on sister taxa suggest single mating: the extremely rapid copulations of *Habropoda pallida* and the apparent absence of a post-insemination courtship signal phase are indicative of a rapid loss of receptivity by females once they have mated[202]. In general, solitary bees are considered monandrous and it was found true for other species from that tribe[213,214].
***Megachile rotundata:*** Based on head capsule width and body weight.
