## supplementary information for "Data-driven analyses of social complexity in bees reveal phenotypic diversification following a major evolutionary transition"

### Table of Contents

**Methods**

**Trait details** ... 3

**Sensitivity analyses** ... 9

**Testing for selection of candidate genes for social complexity** ... 9

**Results**

**Dimensionality reduction** ... 12

**The evolutionary history of social complexity** ... 16

**Sensitivity analyses** ... 20

**Testing for selection of candidate genes for social complexity** ... 23

**References** ... 28

### Methods

#### Traits details

**Table S1.** Socially related traits used in our dataset. We examined relevant papers in *Google Scholar* using the "Search words" for each focal species name. For each trait, we provide the literature supporting the significance of the trait to sociality (numbers in "Social trait"). Ecological contexts include *natural* – a natural colony in a wild\urban area or natural colony taken into a field observation hive; *field*- colonies freely foraging in the field (founded artificially or by using a trap nest box); *free foraging* - lab colony with access to forage freely (semi-natural); *lab* – a colony in complete lab conditions with artificial feeding.

| **Social trait** | **Definition and ecological context** | **Search words** | **Notes** |
|---|---|---|---|
| Colony foundation [1–6] | The number of bees participating in the swarming\absconding or the founding of the colony; The minimal number of bees needed to establish a new colony. Due to high variance and the difficulty to assess accurately thousands of bees, it was log10 transformed.<br>Ecological context: natural; field. | "swarming" OR "colony reproduction*" OR "colony establishment*" | In Meliponini, the swarming can be divided into several phases, each with a different number of workers (with or without gynes). In this case, we took the total number of bees that occupied the nest at the beginning and included all phases. If there was no data available on the number of bees in swarming, we took the minimal colony size reported as an estimation, as long as it was plausible based on actual swarming data of closely related Meliponini taxa (i.e., 10%-40% of the mother-nest, or less than 300 individuals[7]). In some facultatively social species, there can be a certain (usually low) proportion of nests found by more than one individual. However, since the minimal number is 1, we took this value. |
| Colony size maximum [1–4] | The maximum number of adult females reported in a colony in favorable conditions. Due to high variance and the difficulty to assess accurately thousands of bees, it was log10 transformed.<br>Ecological context: natural; field. | "colony size*" OR "number of adults*" | Estimates can be obtained by counting\estimating individuals or by the number of cells or size of the brood area. |
| Colony size average [1–4] | The average number of adults in a colony in favorable conditions.<br>Due to high variance and the difficulty to assess accurately thousands of bees, it was log10 transformed.<br>Ecological context: natural; field. | "colony size*" OR "number of adults*" | Estimates can be obtained by counting\estimating individuals or by the number of cells or size of the brood area. |

| | | | |
|---|---|---|---|
| Colony longevity [8–10] | Estimated average lifespan of a colony after establishment (years). Ecological context: natural; field. | "colony longevity*" OR "colony life span*" OR "colony survival*" OR "colony mortality*" | In some species, it can be a non-trivial task to define colony longevity. Here, we refer to the nest itself that is occupied consecutively (the queen can be replaced). A nest may be sustained for an extended period, but the colony is not, i.e., when the colony dies and is replaced/raided by a different colony of the same species. In this case, knowing the actual sequence of events is almost impossible. Extreme or suspicious values were excluded or mentioned accordingly; in species that exhibit a solitary phase and a social phase, we refer to the social phase only (i.e., when the daughters of the mother emerge or when two or more individuals are cohabiting in the nest for more than a few days after emergence and showing activity of provisioning or foraging). In other words, the longevity of a colony starts when two or more adults start to oviposit and provision brood until all individuals disperse and as long as a sequence of adults remains (at no point do all residents leave\die). In Bombus - if only data from the foundation phase was available, we inferred five weeks as the estimated time for the emergence of the first workers[11]. When there is no overlap of generations, the colony longevity is 0. |
| Queen longevity [12–15] | Maximum lifespan of a queen from emergence to death (years). Ecological context: natural; field; free-foraging. | "queen longevity *" OR "queen lifespan" OR "queen lived*" | For species with no distinct queens (i.e., Allodapini, Halictidae, Euglossini), we took the maximum value of longevity because, in most cases, there was no discrimination between reproductives\non-reproductives. In addition, we assumed a relatively similar lifespan between females of such simple societies as shown in some Halictidae (Yanega, 1988). Also, in facultatively social species, there was often no discrimination between social\solitary status regarding female longevity, but studies suggest that the activity duration of nests is similar[16,17]. |
| Worker size variation [10,18–23] | Coefficient of variation in mean sizes (mm) of workers taken for head/thorax width, unless mentioned otherwise. Calculated as the standard deviation/mean*100. Ecological context: natural; field; free-foraging. | "head width*" OR "thorax width*" | In most papers, body size was recorded during a limited period that may not cover the full seasonal variation in body size. Thus, the reported coefficient of variation may be an underestimate for some species. For species with no clear worker and queen phenotypes (i.e., Allodapini, Halictidae, Euglossini), the data usually represent the entire female population. |

| | | | |
|---|---|---|---|
| Queen-worker size ratio [10,19–21,23–26] | The ratio of the head or thorax width of queen\worker.<br>Ecological context: natural; field; free-foraging; lab. | "head width*" OR "thorax width*" | For species with no distinct queen–worker phenotypes (i.e., Allodapini, Halictidae, Euglossini), the queen was recognized (in the original paper) based on one or more of the following attributes: ovary size, egg laying, and dominant behavior. In articles with no indication of size differences between dominant and subordinate, we assumed a ratio of 1. This assumption was established after noticing that even in studies that mentioned size differences, these were negligible for our analyses[27–29] (~0.02); For Meliponini, we used data for the Intertegular span, which has been proven to be a good indicator of body size in Bombus[30]. |
| Queen-worker differentiation [25,26,31] | The degree of differentiation between queens and workers (or dominant and subordinate)<br>0-no differentiation<br>1-behavioral differentiation<br>2-physiological differentiation (i.e., post-embryonic ovary size)<br>3-size differentiation<br>4-morphological permanent differentiation<br>Ecological context: natural; field; free-foraging; lab. | "female differentiation*" OR "queen worker differentiation*" | |
| % of worker produced males [32–35] | The proportion of male eggs laid by the workers/subordinates in the reproductive phase.<br>Ecological context: natural; field; free-foraging; lab. | "worker produced males*" OR "eggs laid by workers*" OR "worker eggs*" OR "production of males*" OR "male production*" OR "reproductive workers*" OR "worker ovaries*" OR "active ovaries*" OR "worker reproduction*" OR "ovarioles number*" OR "number of ovarioles*" | For few Xylocopinae species these data can be an underestimate because a female can become reproductive only in the second year after emergence; Unless mentioned otherwise in the original article, species with no reproductive skew (i.e., solitary, subsocial, communal), were given values of 100 (i.e., each female lays all eggs for herself); For Bombus, these data were calculated during the competition phase (i.e., worker reproduction phase). |
| Queen fecundity [36,37] | Number of eggs laid per day by a queen. We included only studies in which the queen was in appropriate conditions during the reproductive season.<br>Ecological context: natural; field; free-foraging; lab. | "queen fecundity*" OR "queen fertility*" OR "eggs per year*" OR "colony growth*" OR "colony development*" OR "egg laying rate*" OR "queen oviposition*" OR "brood production*" | In Bombus – it is calculated during the linear growth phase; The lower boundary for eggs laid per day is 1. Thus, for species that lay an egg every few days, we used the value 1. |

| | | | |
|---|---|---|---|
| Worker/queen ratio for ovarioles number [38,39] | Ratio of ovarioles number in ovaries between worker and queen in queenright colonies; (worker/queen).<br>Ecological context: natural; field; free-foraging; lab. | "worker produced males*" OR "eggs laid by workers*" OR "worker eggs*" OR "production of males*" OR "male production*" OR "reproductive workers*" OR "worker ovaries*" OR "active ovaries*" OR "worker reproduction*" OR "ovarioles number*" OR "number of ovarioles*" | In Meliponini - if there was variation among queens (miniature and regular sized), we took the average; Completely sterile workers (occurring only in Meliponini) were given a value of 0. |
| % workers mated [40,41] | The percentage of mated workers in a colony.<br>Ecological context: natural; field; free-foraging; lab. | "worker produced males*" OR "eggs laid by workers*" OR "worker eggs*" OR "production of males*" OR "male production*" OR "reproductive workers*" OR "worker ovaries*" OR "active ovaries*" OR "worker reproduction*" OR "ovarioles number*" OR "number of ovarioles*" | |
| Sex ratio [32,42–45] | Gyne to male ratio during the pupa stage or shortly after emergence from the pupa; calculated as (gynes\(gynes+males)). Unless mentioned otherwise data were collected during the entire year\reproductive phase.<br>Ecological context: natural; field; free-foraging; lab. | "sex ratio*" OR "percent of gynes*" OR "percent of males*" OR "production of reproductives*" OR "queen production*" OR "gyne production*" OR "male production*" OR "production of sexuals*" | We aimed to include data collected over multiple years. For species with no distinct queen and worker phenotypes, we considered all females as reproductive. This index may vary between years, seasons, populations, and colonies. |
| Relatedness [40,46,47] | Genetic relatedness between workers\females in a colony.<br>Ecological context: natural; field. | "relatedness" OR "mating frequency*" | In species without a clear worker caste, the relatedness refers to all females in the nest. |

| | | | |
|---|---|---|---|
| Feeding type [48–52] | Feeding type of the larvae<br>1 - mass provisioning by the queen<br>2 - mass provisioning by the queen and workers<br>3 - mass provisioning by workers<br>4 - progressive provisioning by the queen<br>5 - progressive provisioning by the queen and workers<br>6 - progressive provisioning by workers<br>Ecological context: natural; field; free-foraging. | feeding* OR provisioning* | In bumblebees, both types of food provision are present: ‘pocket-makers’ were classified as mass provisioners and ‘pollen-storers’ as progressive provisioners. |
| Nest complexity [53–56] | Number of structure types in a nest; structure types: brood rearing (in any shape, such as comb, cluster, etc.), entrance, food storage, nest envelope/ involucrum, nest walls/ batumen, burrow, draining duct; facultatively occurring structures got a value of 0.5.<br>Ecological context: natural. | "nest architecture*" OR "involucrum*" | A structure must be built by at least two individuals; The same structure for the same purpose was considered as one (e.g., many identical combs for brood rearing); Stingless bee researchers occasionally use “propolis” (or “geopropolis”) as a synonym for cerumen or an equivalent of “batumen”. |
| Overlap of generations [10,24] | Overlap of generations;<br>0-no overlap<br>0.5-overlap between mother and developing brood<br>1-complete overlap between adults<br>Ecological context: natural; field. | Natural history | |

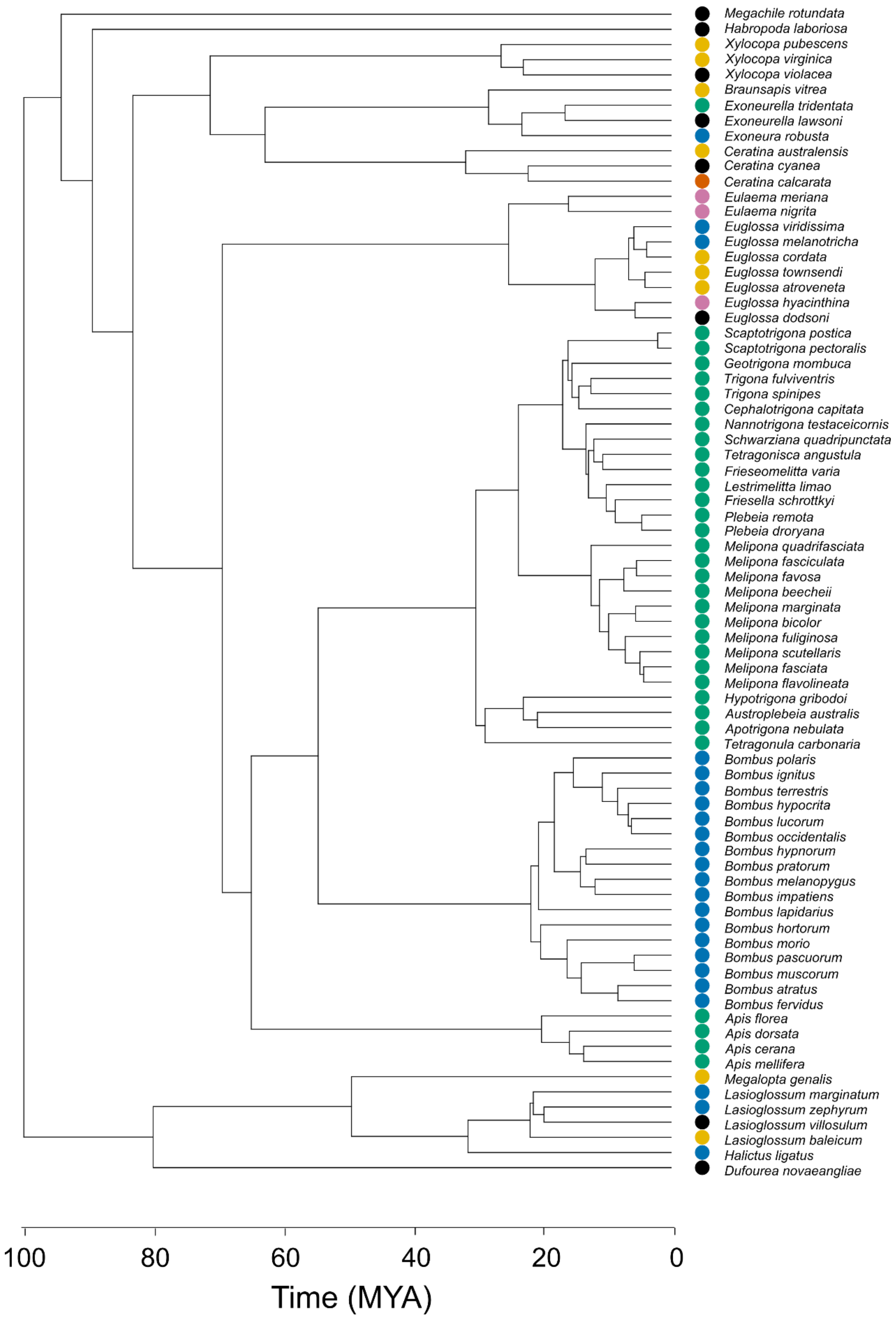


**Figure S1.** The phylogenetic tree presented in Fig. 1b, including species names. Colors show the common social ladder classification of each species as in the main text. Modified from Henríquez-Piskulich et al.[57].

**Sensitivity analysis**

To evaluate the robustness of our results, we conducted a series of sensitivity analyses. These tests were conducted to verify whether our qualitative conclusions remained consistent under various assumptions and data structures.

Removal of traits

First, in order to assess whether specific traits bias our results, we reanalyzed the data after excluding each trait individually. We performed 17 separate PCA analyses and generated a single PCA plot showing the 95% confidence interval ellipse for each species, calculated based on the mean and standard deviation from all 17 analyses. To generate a single plot that remains coherent across multiple analyses, we preserved the orientation of the original PCA axes (i.e., multiplying the axes by -1) in cases of axis rotation.

Missing data threshold and taxonomic bias

Our dataset includes 15% missing data, and the number of missing data entries ranged from 0-47% per species. To assess the influence of missing data on our conclusions, we repeated our analyses using subsets of the data that included only 10%, 8%, and 6% total missing data. Applying these thresholds resulted in the removal of 14, 21, and 31 species from the original dataset, respectively. These analyses offered additional insights into the potential effects of taxonomic sampling bias in our dataset because the species most frequently removed due to missing data were primarily stingless bees and bumble bees.

**Testing for selection of candidate genes for social complexity**

To investigate the potential association between our phenotypic space of social complexity with genomic signals of selection we used selection tests based on the non-synonymous to synonymous substitution ratio (dN\dS). The analysis was based on the availability of genomic data and varied between 10-18 species for each gene. We focused on six candidate genes previously associated with various functions related to social complexity (e.g., reproductive bias, dominance, brood care, etc.). Orthologs for 12 species were taken from Hymenoptera Genome Database[58]. Following the pipeline of Jeffares[59], the remaining orthologs were identified using a BLAST search (using the tblastn feature) on the NCBI database for each species, with *Apis mellifera*'s well-annotated genome serving as the reference (using an e-value cutoff of $10^{-5}$). Reciprocal BLAST hits were treated as tentative ortholog predictions (for limitations see [60]). Following Yang[61], we used the coding domain sequence (CDS) for all the tested genes. We used the MUSCLE codon-based alignment algorithm as implemented in MEGA version 11.0.11 for multiple sequence alignments[62]. Sequence alignments of all genes were carefully reviewed and trimmed in cases of poor alignment, and any sequence gaps were removed. For each of the focal genes, we generated a species gene tree using maximum likelihood as implemented in MEGA[62].

To test for a signature of positive selection we used codeml of the PAML 3.14 software[63] with the EasyCodeML wrapper[64]. In codon analysis, the dN\dS ($\omega$) value can be <1, =1, or >1, indicating purifying (negative) selection, neutral evolution, and positive selection, respectively. We used the branch model, which detects positive selection acting on particular branches of the tree, allowing us to categorize species (i.e., different branches) into different categories and to compare a selected number of species against the rest[63]. In other words, we compared the selection pattern (dN/dS) in focal species relative to the background selection. The categories we used were based on taxonomic lineage, social ladder categories, and the social phenotype groups (A-D) in our analyses. Each category was evaluated using a likelihood ratio test (LRT) against a null model, where all branches on the tree are assumed to have the same dN/dS rate. A Bonferroni correction was applied for multiple comparisons, with $\alpha$=0.05. We then assessed the LRT values across all categories to identify the most supported one (i.e., with the highest log likelihood). When performing the test, we took several precautionary measures. First, to avoid incorrect comparisons (e.g., caused by neofunctionalization or subfunctionalization), we only used single-copy genes, following Shell et al.[65]. Second, to increase the possibility of detecting a signal as suggested by Anisimova[66], we used genes with medium-large sequence length (260 codons or more). For each gene, we report the transition/transversion ratio between homologous strands of DNA (ts\tv). To provide information regarding the structure of the sequences and their degree of divergence, Table S2 also includes details of the of synonymous and nonsynonymous substitution rates, dS and dN respectively.

We then examined the association between the PCA values (Fig. S4) with the selection signals we identified in the six candidate genes. We calculated the pairwise dN/dS of all species with the solitary *Megachile rotundata*. We then used these values to perform a linear regression analysis of the dN/dS values with PCs 1-4. Five out of the 21 species for which we had genetic data but were not included in our phenotypic analysis, were replaced with the most socially and phylogenetically similar species from our dataset (Table S3).

**Table S2.** Details of the genes and sequences used to measure dN/dS.

| Gene | GeneID | kappa (ts/tv) | length dN | length dS | NCBI |
|---|---|---|---|---|---|
| *dunce* | 411288 | 3.27485 | 0.187 | 8.6667 | https://www.ncbi.nlm.nih.gov/gene/411288 |
| *TOR* | 409393 | 4.192 | 0.123 | 5.987 | https://www.ncbi.nlm.nih.gov/gene/409393 |
| *syx1a* | 410279 | 2.51021 | 0.0541 | 2.4356 | https://www.ncbi.nlm.nih.gov/gene/410279 |
| *IRS* | 408438 | 4.13826 | 0.4276 | 4.3793 | https://www.ncbi.nlm.nih.gov/gene/408438 |
| *InR-2* | 725827 | 2.23243 | 0.3523 | 7.8231 | https://www.ncbi.nlm.nih.gov/gene/725827 |
| *InR* | 411297 | 2.42854 | 0.3121 | 9.9717 | https://www.ncbi.nlm.nih.gov/gene/411297 |

**Table S3.** Species included in the dN/dS analyses, with the categories used for each species in the branch model. 'v' marks the data availability of species for each gene in the analysis.

| species | sociality | grp | phylogeny | dnc | syx | IRS | TOR | InR-2 | InR |
|---|---|---|---|---|---|---|---|---|---|
| *Apis mellifera* | advanced eusocial | D | Apinae | v | v | v | v | v | v |
| *Apis florea* | advanced eusocial | D | Apinae | v | v | v | v | | v |
| *Frieseomelitta varia* | advanced eusocial | D | Apinae | v | v | v | v | | |
| *Melipona quadrifasciata* | advanced eusocial | D | Apinae | v | v | v | v | v | v |
| *Tetragonula carbonaria* | advanced eusocial | D | Apinae | v | | v | v | | |
| *Exoneurella tridentata* | advanced eusocial | E | Xylocopinae | v | v | v | v | v | v |
| *Heterotrigona itama*[a] | advanced eusocial | D | Apinae | v | | | | | |
| *Megalopta genalis* | parasocial | B | Halictidae | v | v | v | v | v | v |
| *Lasioglossum albipes*[b] | parasocial | B | Halictidae | v | v | v | v | v | v |
| *Eufriesea Mexicana*[c] | parasocial | B | Apinae | v | v | v | v | v | v |
| *Ceratina australensis* | parasocial | B | Xylocopinae | | | v | | | |
| *Bombus terrestris* | primitively eusocial | C | Apinae | v | v | v | v | | v |
| *Bombus impatiens* | primitively eusocial | C | Apinae | v | v | v | v | v | |
| *Exoneura robusta* | primitively eusocial | B | Xylocopinae | v | v | v | v | | |
| *Euglossa dilemma*[d] | primitively eusocial | B | Apinae | | v | v | v | | |
| *Megachile rotundata* | solitary | A | none | v | v | v | v | v | v |
| *Dufourea novaeangliae* | solitary | A | Halictidae | v | v | v | v | v | v |
| *Habropoda laboriosa* | solitary | A | none | v | v | v | v | | |
| *ceratina japonica*[e] | solitary | A | Xylocopinae | | | | v | | |
| *Ceratina calcarata* | subsocial | A | Xylocopinae | v | v | v | v | v | v |

letters denote species that were not included in the social trait dataset and were therefore replaced in the dN/dS analysis with phylogenetically related taxa with assumingly a similar level of social complexity. *a- Trigona fulviventris; b- Lasioglossum baleicum; c- Euglossa hyacinthine; d- Euglossa viridissima; e- Ceratina cyanea.*

### Results

##### Dimensionality reduction

The broken stick method indicated four statistically significant PCs which together explain ~70% of the variation. For each PC, a trait was considered to make a significant contribution if it accounted for more than 6% of the variation, which is more than would be expected by chance (100 divided by 17 traits). Out of 17 traits included in our dataset, eight traits contributed significantly to the variation of PC1, whereas PCs 2, 3, and 4 each had 4-6 significant contributing traits (Fig. S3). Our results emphasize that a single major component (i.e., PC1) only partially explains the phenotypic variation between species and the relationship between social complexity traits, as indicated by the mixture of traits in the PCs. This means that using a single trait or combination of traits as predictors for the social complexity phenotype of bees can be misleading

Integration of the patterns emerging from comparing all possible pairs of the four significant PCs (Fig. S4) supports a data-driven phenotypic clustering into four groups termed A-D (Fig. 2). For example, species that are commonly classified as solitary, communal, and subsocial were clustered together, and this cluster was positioned close to another cluster containing parasocial and non-*Bombus* primitively eusocial species. Similarly, bumble bees, honey bees, and stingless bees were often clustered in proximity. However, along each PC the species are partitioned differently, and areas display a varying degree of overlap. In PC1 we observe a pattern of increased values partially corresponding to the commonly used social classification from solitary to primitively eusocial to advanced eusocial. In PCs 2, 3, and 4, the relationships between species are more complex, with no clear linear gradient from one social classification to another. PC3, which is dominated by the ovarioles ratio and relatedness traits, clearly distinguishes between the honey bees and the stingless bee *Frieseomelitta varia* on one side, and the rest of the species on the other. In PC4, which was mainly affected by colony longevity, queen longevity, and queen-worker size ratio, we observe a continuum from bumble bees to stingless bees. These contrasting patterns highlight the importance of considering the multidimensional nature of social complexity in bees and that ignoring this structure might lead to an oversimplified interpretation of social evolution.

The analysis of PCA and UMAP without a phylogenetic correction provided identical results in the UMAP and similar results in the PCA. The main difference observed is a clearer separation between bumble bees and stingless bees, and a higher proportion of variation (53%) explained by PC1, which again suggests that phylogeny plays a critical role in the social phenotypes of bees.

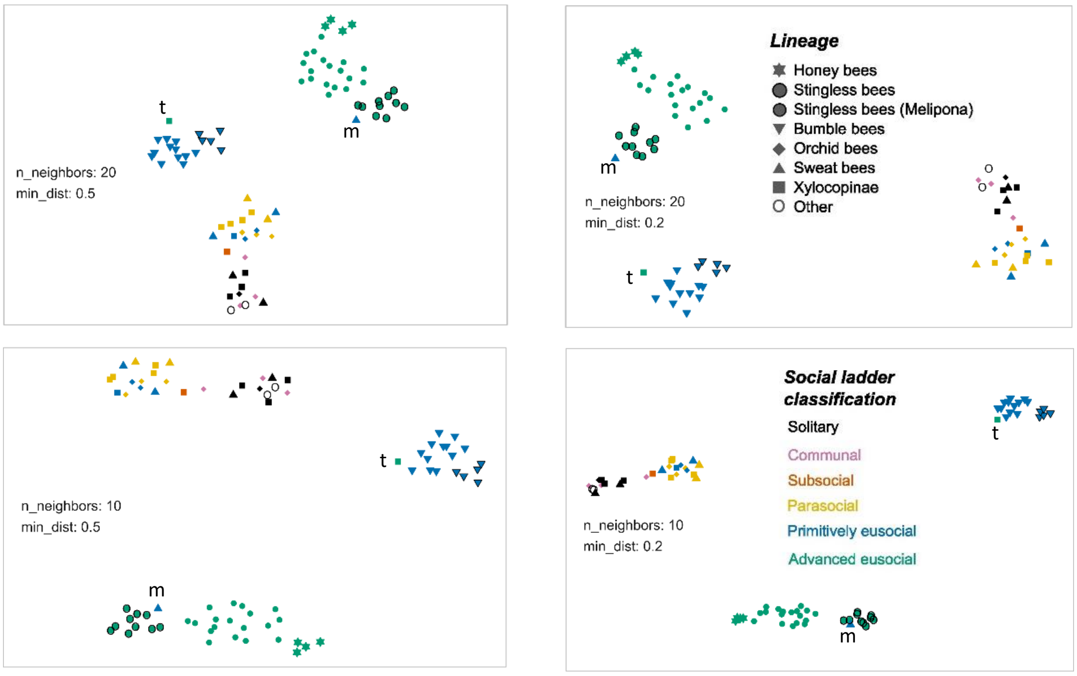


**Figure S2**: UMAP analysis with different clustering parameter values for the number of neighbors and minimum distance (n_neighbors and min_dist, respectively). Legend color and shape as in Fig. 2. Upside-down triangles with or without a black outline refer to bumble bees with mass (pocket makers), or progressive (pollen storers) provisioning, respectively. *E. tridentata* and *L. marginatum* are marked with 't' (green square) and 'm' (blue triangle), respectively.

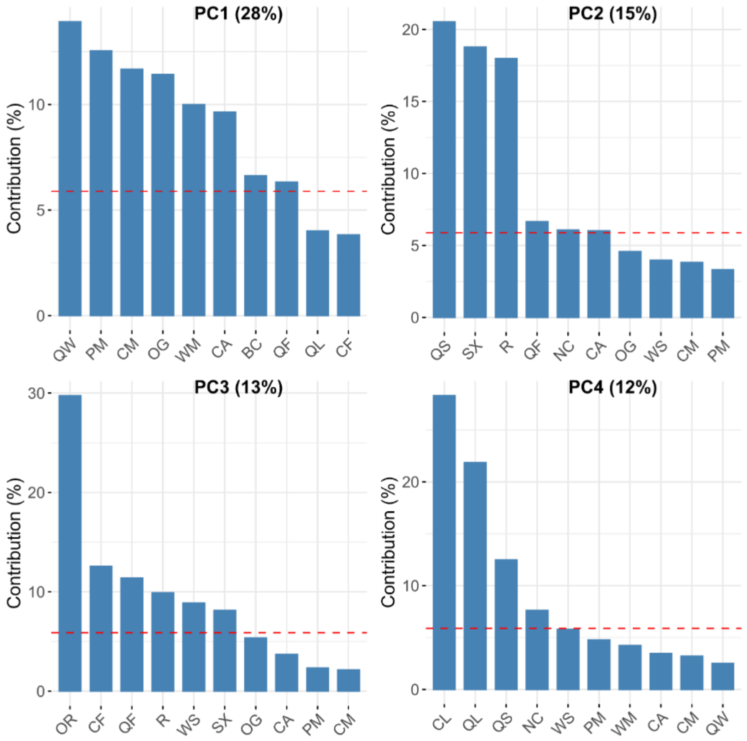


**Figure S3.** Contribution of traits to the four main PCs in our PCA. The variance explained by each component is reported in parenthesis next to the PC number. Red reference line corresponds to the expected value if the contribution of all traits was uniform.

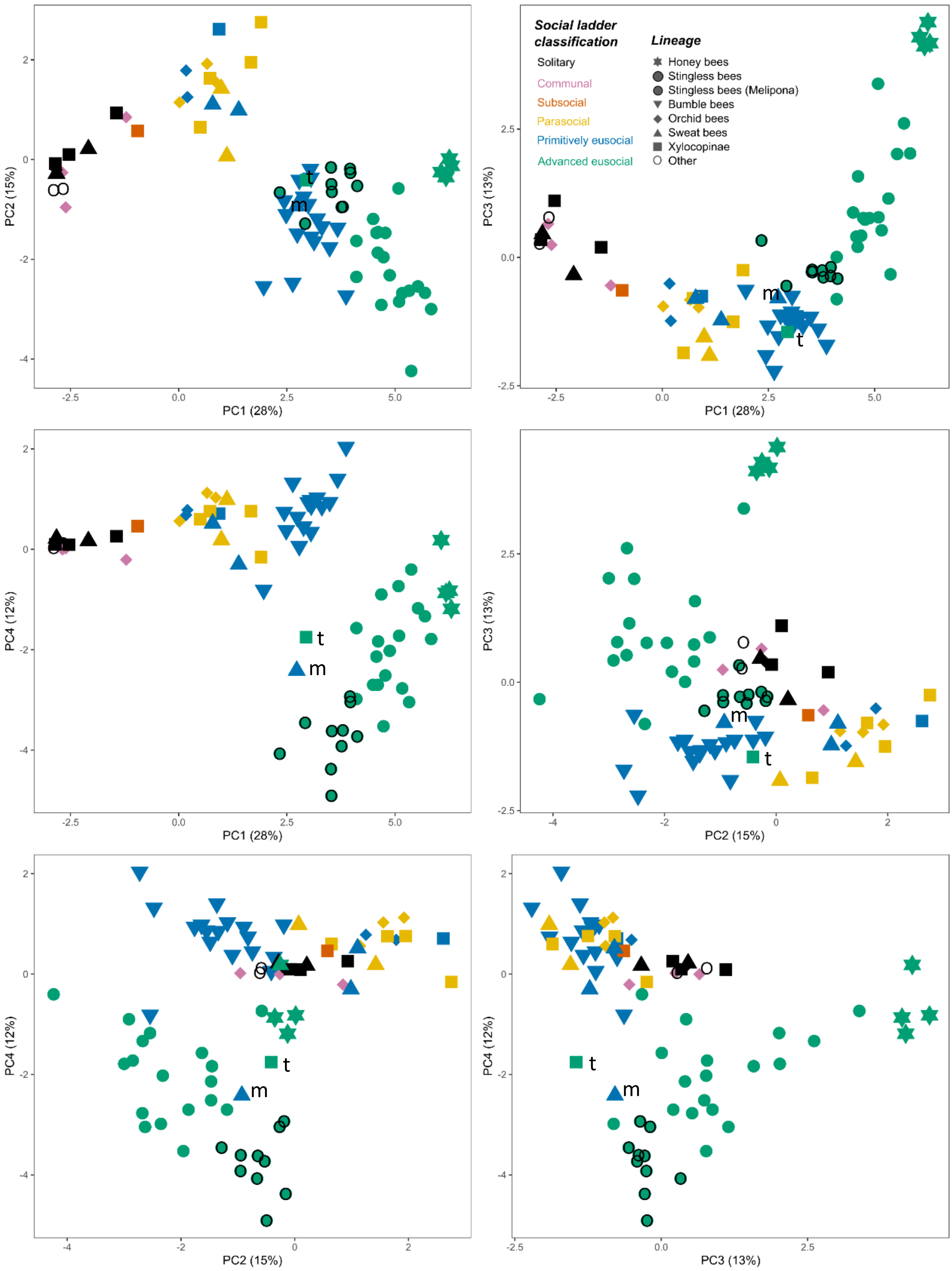


**Figure S4.** Combinations of the four main PCs. Colors correspond to the social ladder classification, and symbol shape refers to the taxonomic clade of the species. Details of plots are as in Fig. S2 above.

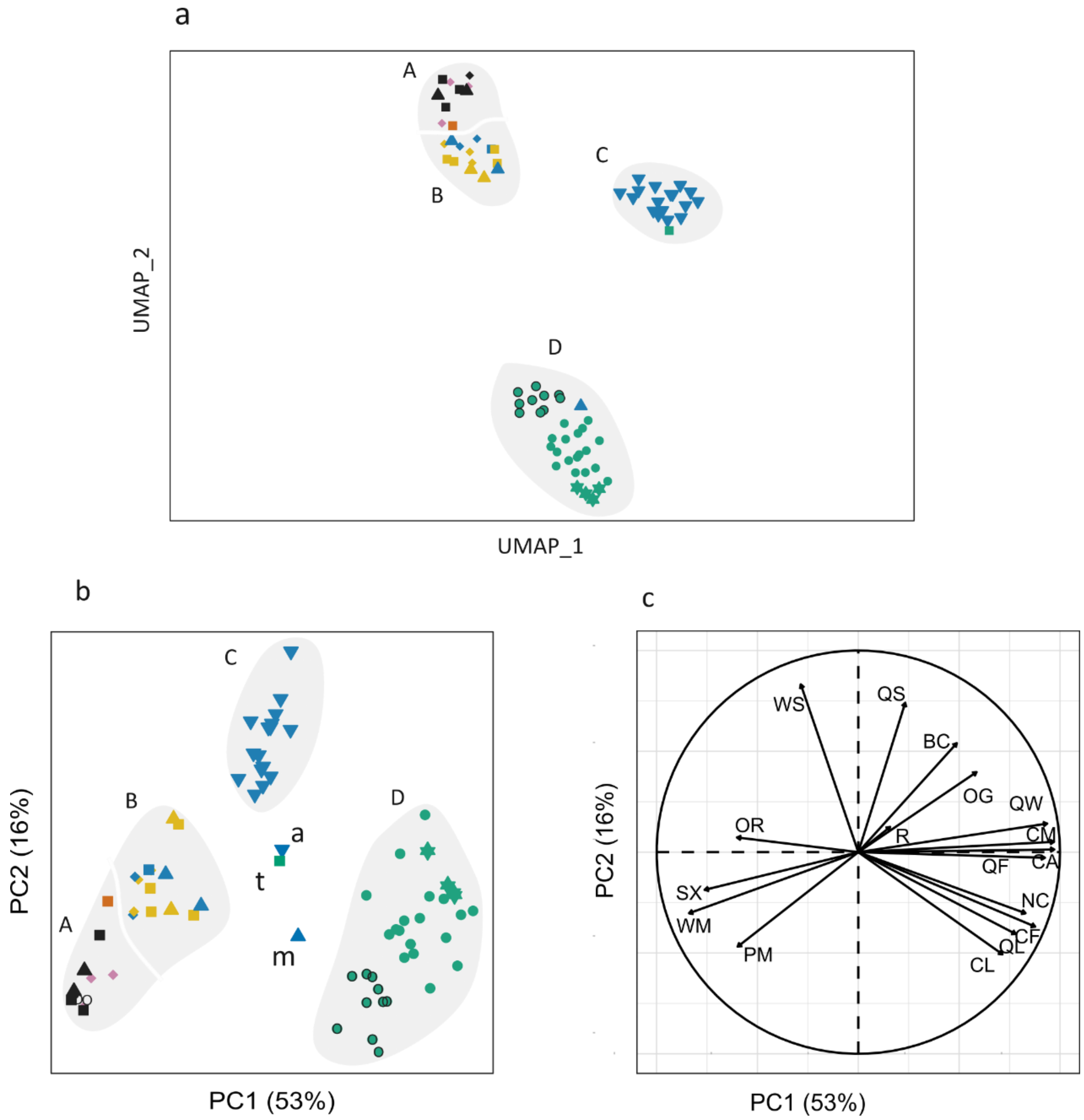


**Figure S5.** PCA and UMAP with no phylogenetic correction. *E. tridentata*, *L. marginatum*, and *B. atratus* are marked with 't', 'm', and 'a', respectively. In the PCA, these species do not cluster with groups A-D and emerge as outliers. Legend as in Fig. S2

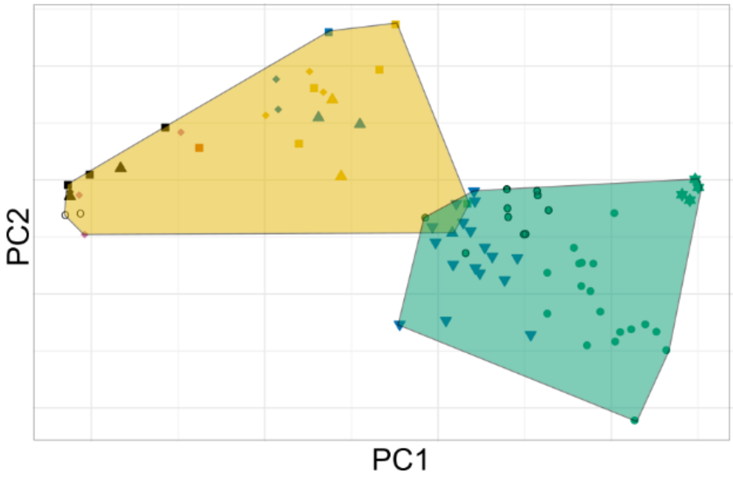


**Figure S6.** The area of the phenotypic space occupied by corbiculate species, which are all defined as eusocial (green polygon), and the remaining species, which range from solitary to primitively eusocial. The plot shows phylogenetically corrected PC1 and PC2 with convex hulls around polygons.

**The evolutionary history of social complexity**

Ancestral state reconstruction on the four significant PCs delineates multiple evolutionary trajectories for increased social complexity in bees (Fig. S7). The trajectories could not be assigned to specific lineages, except for corbiculate bees in PC1, honey bees in PC3, and stingless bees in PC4. High values of PC1 appear irreversible in the corbiculate, with no evidence for a reversion toward values lower than that of the ancestor of corbiculate bees. However, there are substantial increases and decreases within the phenotypic spaces occupied by honey bees, stingless bees, and bumble bees. This pattern is consistent with evolutionary divergence (Fig. 5).

The ancestral state reconstruction of PC3, which is dominated by the ovarioles ratio (a physiological measure for the reproductive skew between queens and workers), suggests a separation of honey bees from the other lineages at approximately 60 million years ago (mya), with the bumble bees more similar to species in groups A and B than to the stingless bees (Fig. S7). Overall, PCA values do not consistently increase across all PCs as we approach the present, supporting the notion that there is no single optimum level of high social complexity to which all lineages are drawn. Moreover, the evolutionary trajectory of social complexity in corbiculate bees did not appear to pass through intermediate levels (as seen in the phenotypes of extant parasocial and primitively eusocial species of sweat bees, Xylocopinae bees, and orchid bees; Fig. S8).

Adaptive regime shift detection analysis for all four PCs (instead of only PC1 and PC2 as shown in the main text) identified four shift positions (Fig. S9). The position of shifts varied slightly between results, but all solutions received the same support with shifts in each clade of honey bees, stingless bees, the *Melipona* genus of stingless bees, and bumble bees.

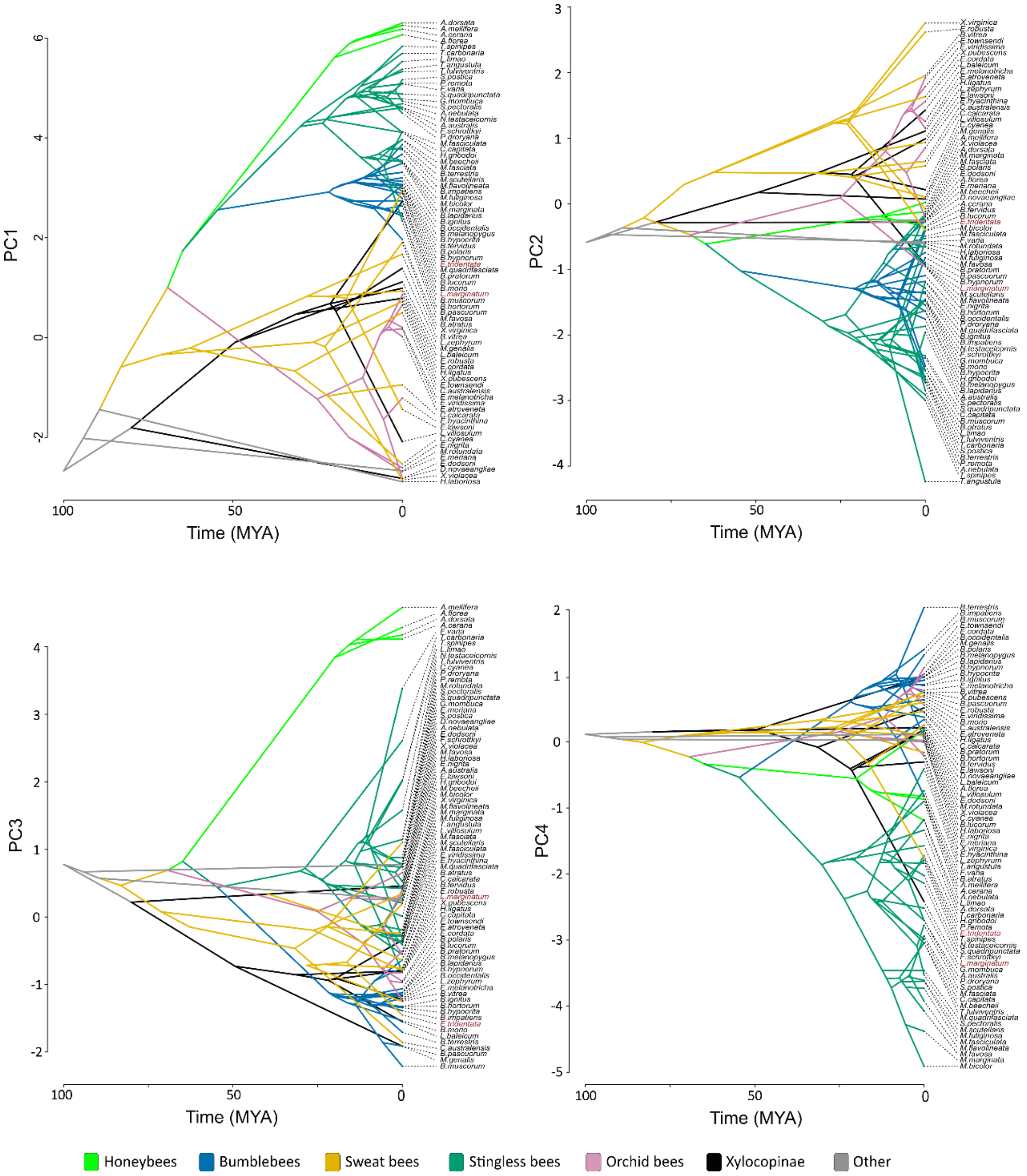


**Figure S7.** Ancestral state reconstruction for the four main PCs in the PCA of social complexity. Colors correspond to the different lineages of bees. *E. tridentata* and *L. marginatum* names are marked with red. Root value was fixed to solitary based on the values of *M. rotundata*. Other details as in Fig. 3a.

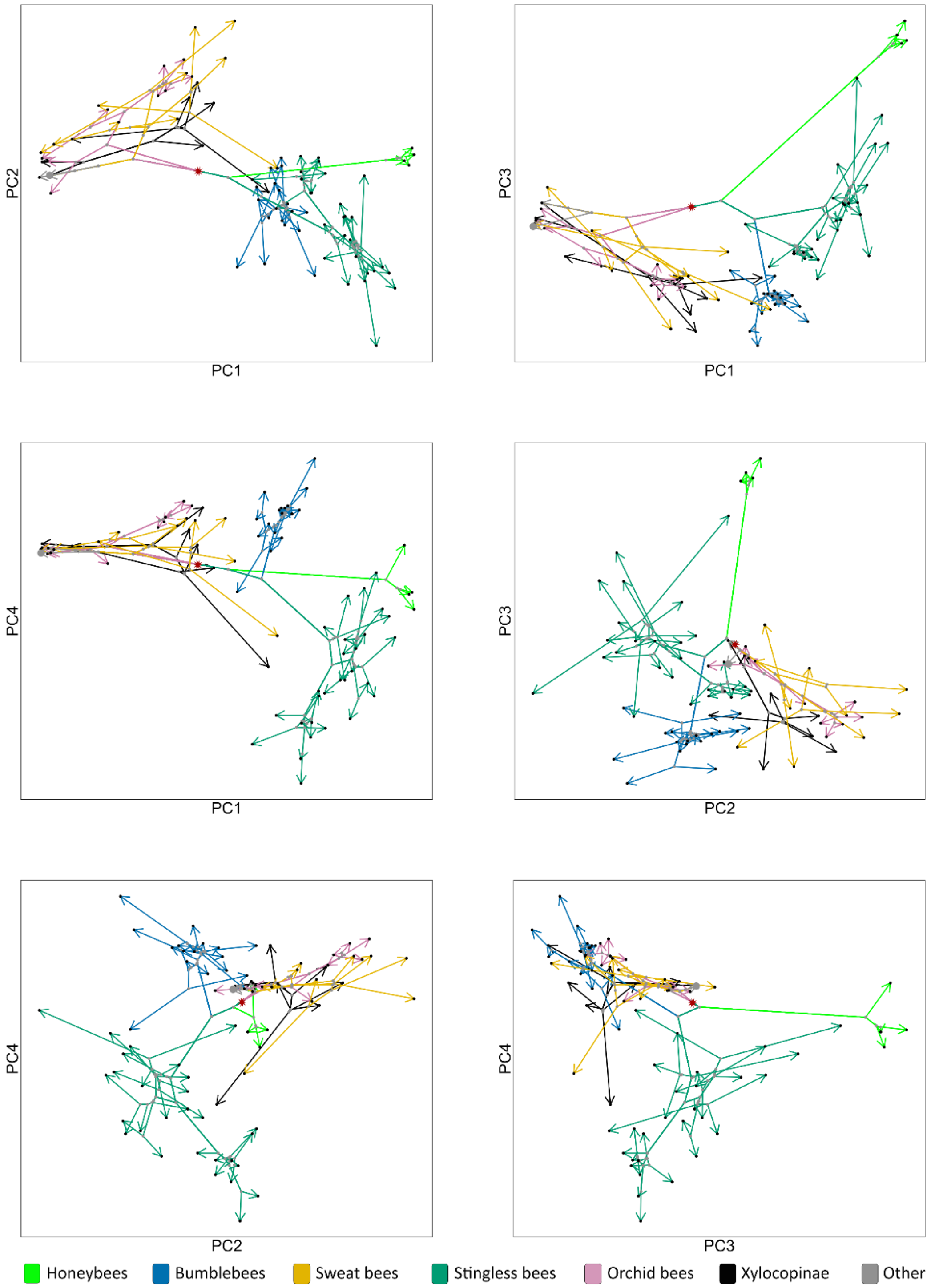


**Figure S8.** Ancestral state reconstruction for each combination of the four significant PCs in the PCA of social complexity. Arrows depict the inferred evolutionary route of species within the phenotypic space presented in each panel. Big gray dots mark the assumed position of the most recent common ancestor of species (MRCA). The assumed ancestor of honey bees, stingless bees, and bumble bees is denoted with a red asterisk. Other details as in Fig 3b.

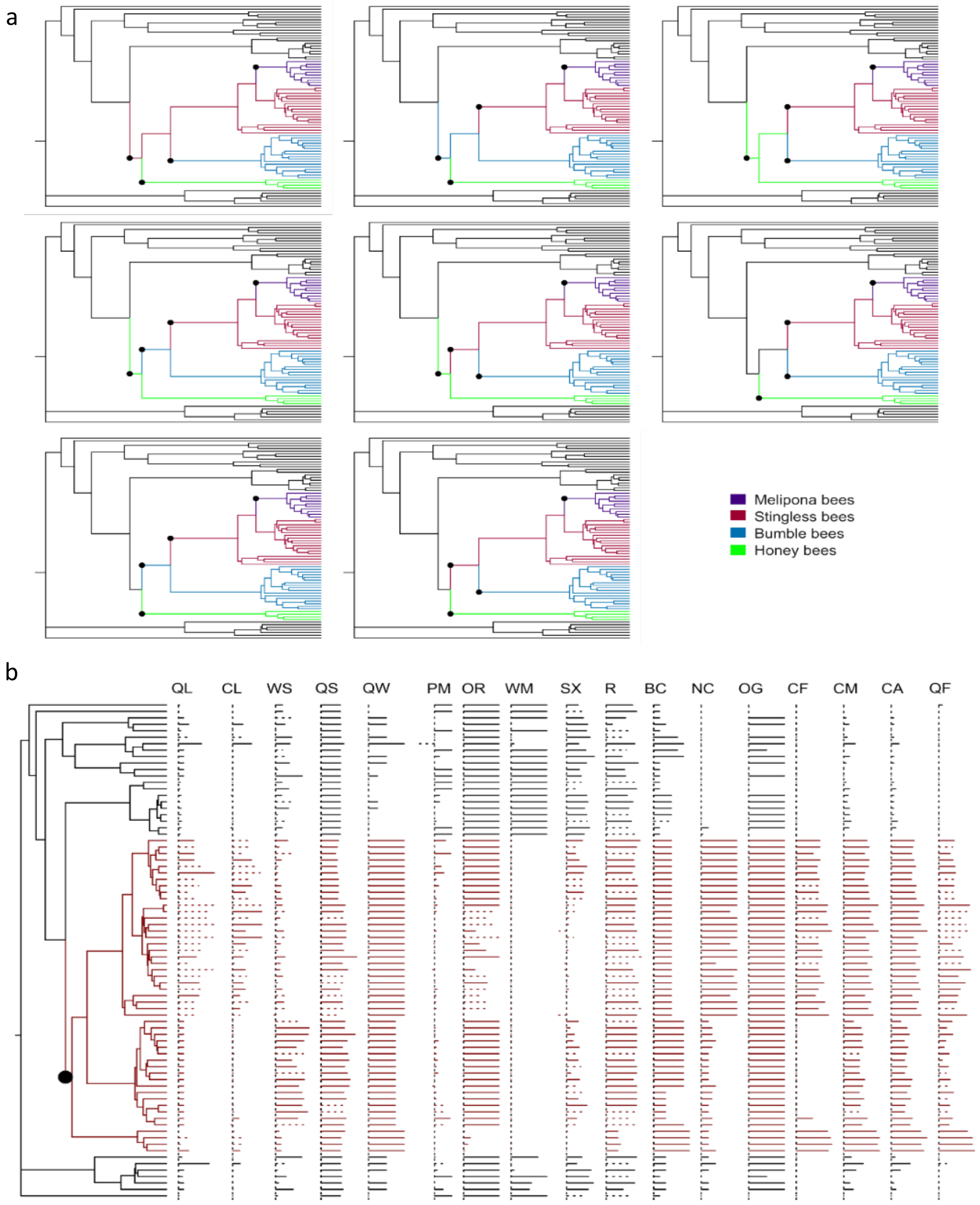


**Figure S9.** Adaptive regime shift detection analysis. **(a)** Multiple equivalent solutions for analyzing the four main PCs together. All solutions indicate four adaptive regime shifts in each clade of honey bees, stingless bees, Melipona genus stingless bees, and bumble bees. **(b)** An analysis using all traits in our dataset. A single regime shift, highlighted by a black circle and red coloring, was detected in the ancestor of honey bees, stingless bees, and bumble bees. Trait values are shown on the right.

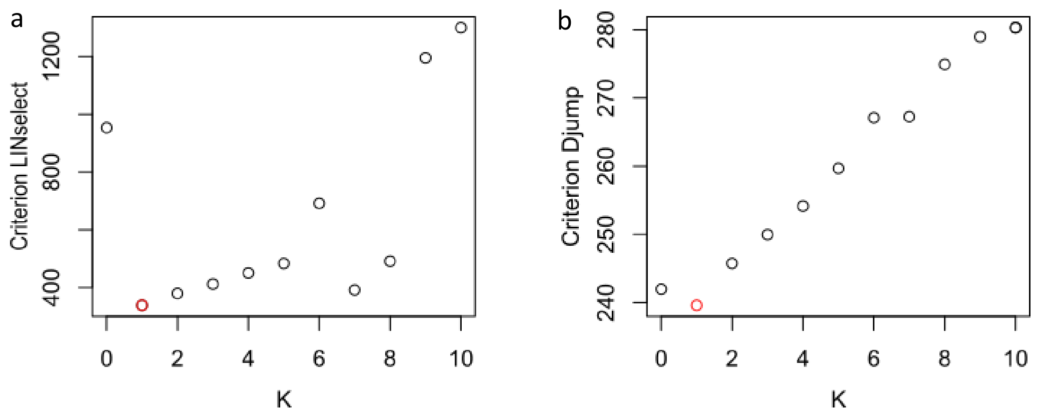


**Figure S10.** Model selection statistic to choose K (number of shifts) in the shift detection analysis, based on PC1 and PC2. Red circles mark the best supported K. **(a)** results based on the 'LINselect' method **(b)** results based on the 'Djump' method.

**Sensitivity analysis**

<u>Removal of traits</u>

The position of the species in the phenotypic space of PC1 and PC2 in our sensitivity analysis of removing traits was similar to the original analysis including all traits (Fig. S11). The confidence ellipses in PC1 and PC2 describe the variability in the social complexity of species considering different sets of traits. The confidence ellipses were small, indicating that species maintained in the broad phenotypic groups (A-D) we describe in the main text. Sensitivity analysis for PCs 3 and 4 remained qualitatively similar, but because fewer traits contribute to the variation in these PCs they are more sensitive to trait removal and showed higher confidence intervals.

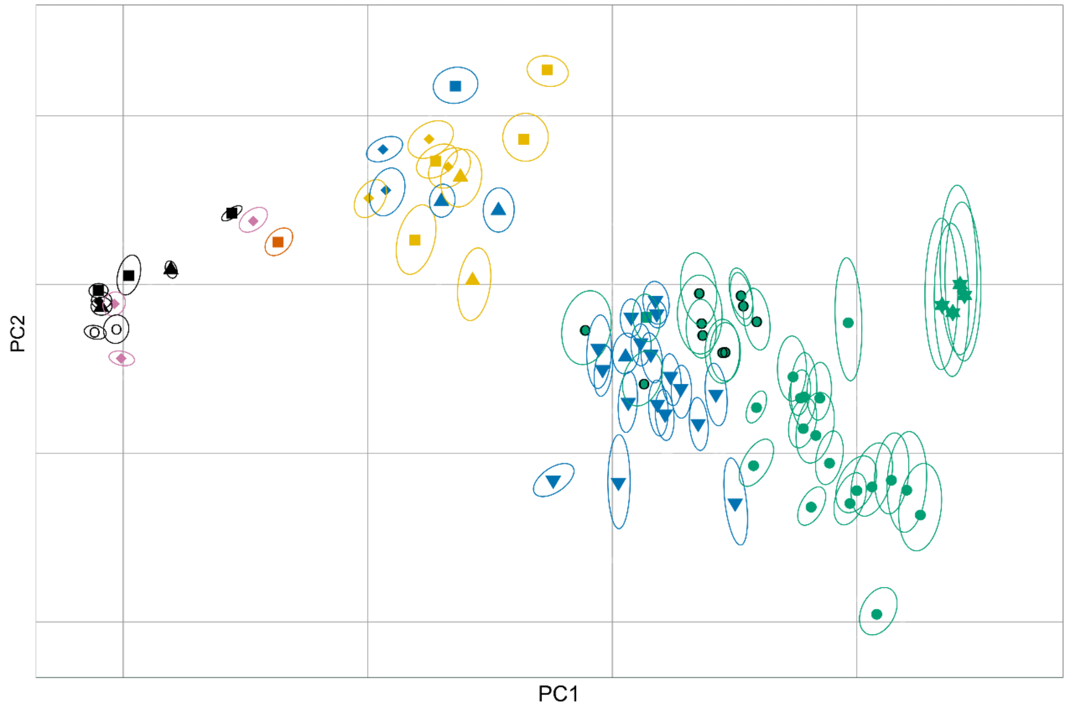


**Figure S11.** Sensitivity analysis for the PCA showing PC1 and PC2. The ellipsoids delineate the 95% confidence intervals for each species, calculated based on the results of 17 analyses, in each one of which we excluded one trait from the PCA analysis.

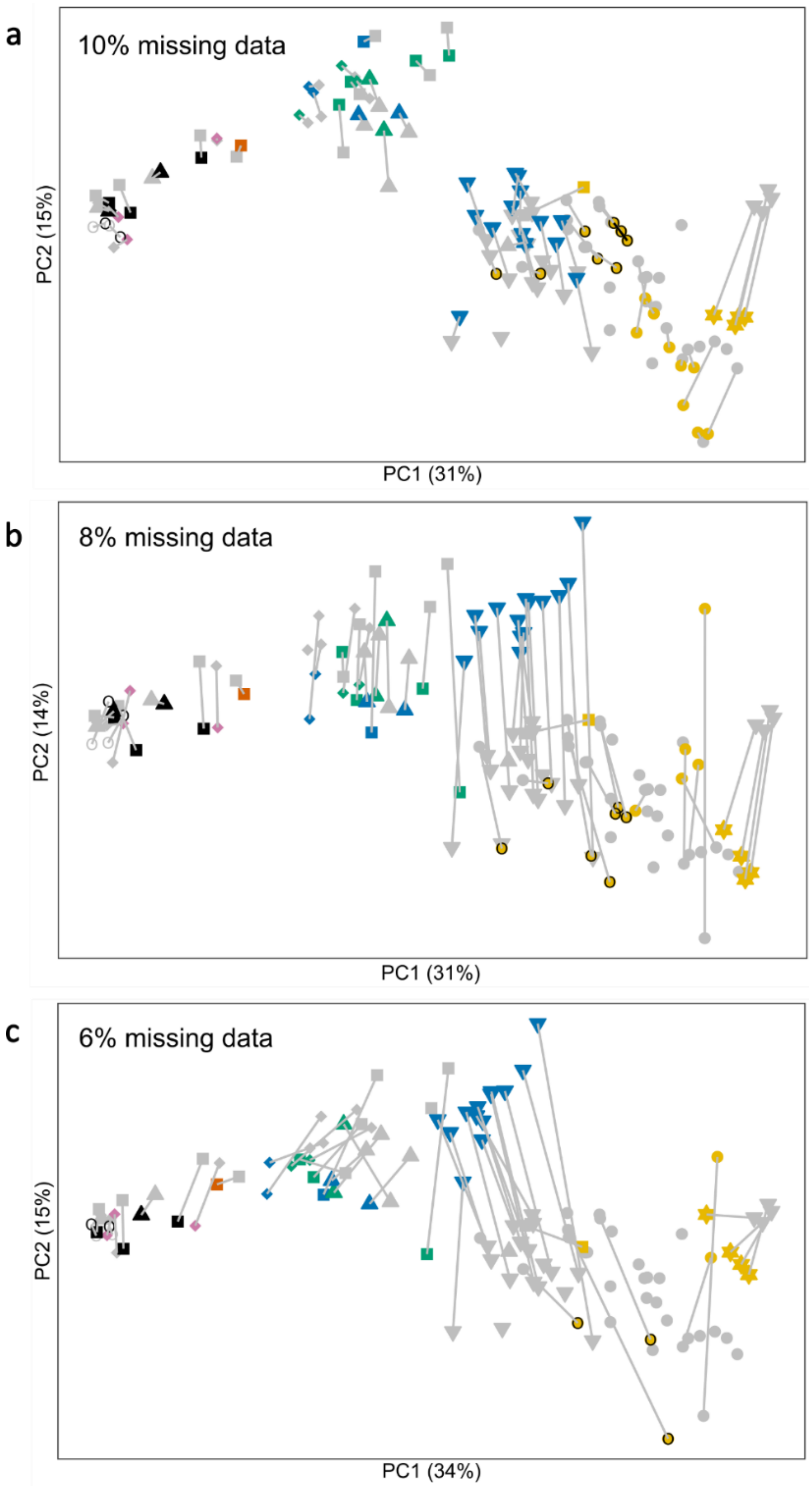


**Figure S12.** Assessing the influence of missing data on our PCA results. The plots show PC1 and PC2 in which we limited the analyses to only species with (a) 10% (63 species), (b) 8% (56 species), and (c) 6% (46 species) missing data, compared to 15% in the analyses presented in the main text with 77 species). Symbol color and shape are the same as in Fig. 2. Gray shapes represent the original position in Fig. 2. The gray lines connect it to the positions of each species in the analysis with the new missing data threshold. Gray shapes with no connecting lines represent the species removed from the analysis due to the change in the missing data threshold.

Missing data threshold

The PCA analysis with a dataset that includes only species with no more than 10% missing data overall preserved the separation between the main phenotypic groups as well as the position of all species, except for a few that changed their position along PC2 (Fig. S11a). PC1 explained 31% of the variation compared to 28% in the original analysis (Fig. S11a). In an analysis where we limited our dataset to include only species with no more than 8% missing data, the number of species was reduced to 56 (Fig. S11b), resulting in more species changing their position, with the changes typically being more extensive (specifically along the PC2 axis). However, the separation based on taxonomy and our social complexity groups remained clear and overall similar. These trends were further exaggerated when the dataset was limited to include only species with no more than 6% missing data (Fig. S11c). Importantly, however, the separation of the main social phenotypic groups (A-D) was evident in all the analyses. Detection of adaptive regime shifts based on PC1 and PC2, using the most stringent criterion of no more than 6% missing data, supported a phenotypic transition in the ancestor of honey bees, stingless bees, and bumble bees (Fig. S13). It also suggested an additional transition in the ancestor of bumble bees, which were indeed further separated from *Melipona* stingless bees compared with the outcomes of the original PCA.

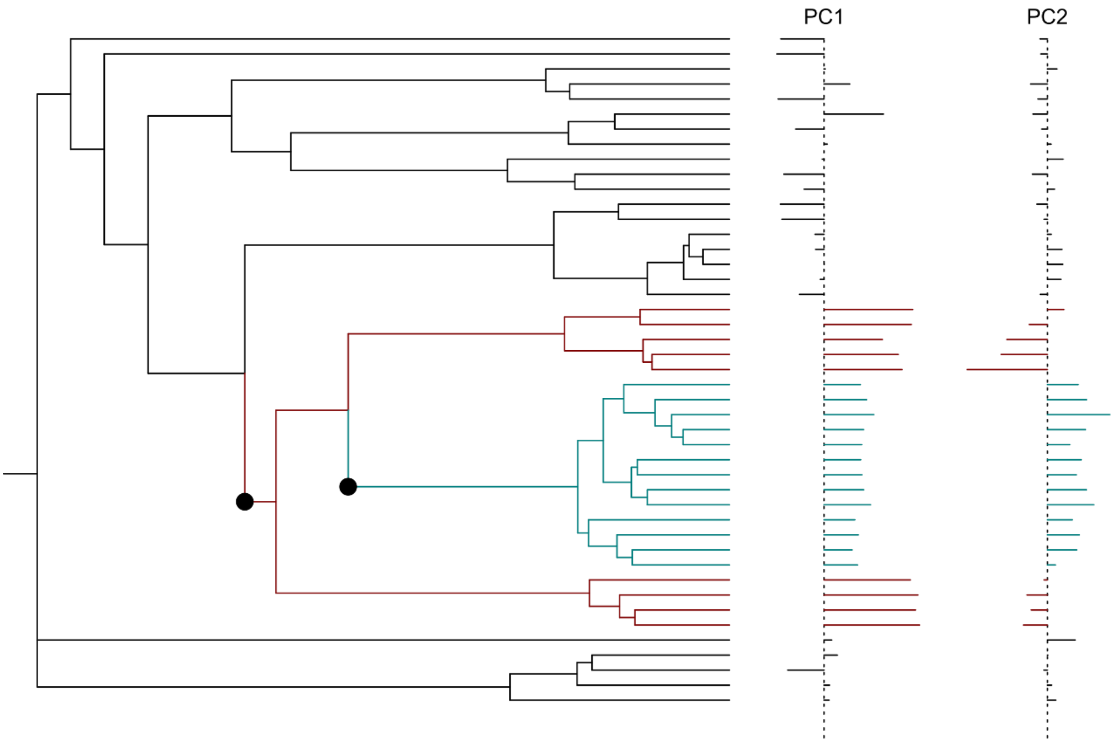


**Figure S13.** Adaptive regime shift detection analysis for a dataset with 6% missing data. Two shifts were detected: in red - in the ancestor of corbiculate bees; in turquoise – bumble bees. Trait values are shown on the right.

**Testing for selection of candidate genes for social complexity**

Out of the six genes we tested, *Dunce* showed the strongest signal for positive selection in bumble bees (relative dN/dS=22.5; Tables S4; S5). *Dunce* is part of a signal transduction pathway involved in learning and memory in Drosophila[68,69] and has emerged as an important gene in the regulation of neural plasticity in both invertebrates and vertebrates[70]. Specifically in social insects, recent studies implicated *dunce*, along with other genes in the cAMP pathway, in social learning[71] and with evidence for rapid evolution in bumble bees[72] and high expression levels of *dunce* in dominant females of the small carpenter bee *Ceratina calcarate* and *Solenopsis invicta* fire ants[73,74]. Selection signatures (pairwise dN/dS values) on *dunce* are positively correlated with PC4 ($R^2$=0.34, $p$=0.01; Tables S6), which is influenced by queen-worker and worker-worker size variation as well as with colony and queen longevity. These results are consistent with selection of *dunce* in bumblebees and suggest that it might be involved particularly in processes associated with size-based division of labor and dominance in bumble bees [22].

*TOR* (target of Rapamycin), which has been repeatedly implicated in the evolution of sociality and specifically in caste differentiation[75–77], showed a low signal of positive selection in a subsocial species *C. calcarata*. *Syx1a*, for which a specific SNP has been associated with social nesting in the facultative social Halictidae *Lasioglossum albipes*[78], showed a low signal of positive selection in branches leading to species from group A in our analyses. Interestingly, *syx1a* also showed a significant negative correlation with PC1 ($R^2$=0.32, $p$=0.02; Table S6), which indeed partitioned species with facultative and obligate social phenotypes. A previous study found a signal for positive selection in social bees in another SNARE encoding gene from the same family, *syntaxin7*[72]. The insulin signaling pathway genes *Insulin receptor substrate 1-B* (*IRS*) and *insulin-like peptide receptor* (*InR*), which have been implicated in caste-related characteristics in ants and bees[75,79,80], showed low signatures of positive selection in *E. tridentata*. Interestingly, we also identified this species as having a recent change in its evolutionary history of social complexity (Fig. S6). The *insulin-like receptor-like (InR-2)* showed a weak signal of purifying selection in bumble bees (Tables. S4 and S5).

**Table S4.** Results for the branch model calculating the dN/dS for six focal genes previously implicated in the evolution of social complexity. '*Function*' - suggested function based on previous studies; '*Best model*' - the best supported branch that showed the highest likelihood compared to a null model where all branches evolve at the same rate; *dN/dS* - the dn/ds ratio for the focal branch relative to the background branches; '*p.value*' - likelihood ratio test *p*-values.

| **Gene** | **Function** | **Best model** | **dN/dS** | ***p*-value** |
|---|---|---|---|---|
| ***Dunce*** | learning and memory | cluster C (bumble bees) | 22.5 | <0.001 |
| ***TOR*** | caste determination\ longevity | subsocial (*C.calcarata*) | 6.1 | <0.001 |
| ***syx1a*** | induce sociality in a facultatively social bee | cluster A | 5.6 | <0.001 |
| ***IRS*** | Caste determination\ brood care | *E. tridentata* | 3.5 | <0.001 |
| ***inR-2*** | Caste determination\ division of labor | cluster C (bumble bees) | 0.15 | <0.001 |
| ***InR*** | Caste determination\ division of labor | *E. tridentata* | 3 | <0.001 |

**Table S5.** Log likelihood results of the branch model of 'codeml' for each gene. $\omega_0$ is the dN\dS of the background branch. $\omega_1$ is the dN\dS of the foreground (selected) branch. *p*-value based on likelihood ratio test with a significance level of α=0.01 adjusted with Bonferroni correction for multiple comparisons. In bold is the best supported model with the highest log-likelihood.

| *Model* | *Log Likelihood* | $\omega_0$ | $\omega_1$ | *p-value* |
|---|---|---|---|---|
| *dunce* | | | | |
| *cluster A* | -5253.9 | 0.036 | 0.004 | 0.00000 |
| *cluster B* | -5272.31 | 0.025 | 0.009 | 0.07060 |
| ***cluster C*** | **-5201.42** | **0.013** | **0.270** | **0.00000*** |
| *cluster D* | -5271.17 | 0.024 | 0.004 | 0.01846 |
| *E. tridentata* | -5273.88 | 0.021 | 0.027 | 0.73813 |
| *solitary* | -5263.76 | 0.030 | 0.003 | 0.00001 |
| *subsocial* | -5267.11 | 0.027 | 0.004 | 0.00022 |
| *parasocial* | -5276.43 | 0.021 | 1.950 | 0.02572 |
| *primitively eusocial* | -5217.92 | 0.012 | 0.185 | 0.00000 |
| *advanced eusocial* | -5269.28 | 0.026 | 0.003 | 0.00227 |
| *Apidae* | -5222.83 | 0.013 | 0.174 | 0.00000 |
| *Halictidae* | -5273.93 | 0.022 | 0.020 | 0.89249 |
| *Xylocopinae* | -5264.28 | 0.030 | 0.005 | 0.00001 |
| *null* | -5273.94 | | | |
| *TOR* | | | | |
| *cluster A* | -32993.36 | 0.022 | 0.016 | 0.00547 |
| *cluster B* | -32996.52 | 0.020 | [] | 0.23591 |
| *cluster C* | -32997.13 | 0.021 | [] | 0.66626 |
| *cluster D* | -32995.55 | 0.021 | [] | 0.06768 |
| *E. tridentata* | -32991.31 | 0.020 | 0.044 | 0.00059 |
| *solitary* | -32995.19 | 0.022 | 0.017 | 0.04397 |
| ***subsocial*** | **-32989.45** | **0.020** | **0.061** | **0.00008*** |
| *parasocial* | -32997.22 | 0.021 | [] | 0.91112 |
| *primitively eusocial* | -32991.28 | 0.020 | 0.035 | 0.00056 |
| *advanced eusocial* | -32996.95 | 0.020 | [] | 0.45718 |
| *Apidae* | -32991.59 | 0.022 | 0.009 | 0.00079 |
| *Halictidae* | -32995.53 | 0.021 | [] | 0.06614 |
| *Xylocopinae* | -32995.16 | 0.020 | 0.027 | 0.04205 |
| *null* | -32997.22 | | | |
| *Syx1a* | | | | |
| ***cluster A*** | **-2792.33** | **0.010** | **0.055** | **0.0000** |
| *cluster B* | -2802.16 | 0.022 | [] | 0.9289 |
| *cluster C* | -2801.2 | 0.023 | [] | 0.1644 |
| *cluster D* | -2798.22 | 0.025 | 0.000 | 0.0050 |
| *E. tridentata* | -2802.07 | 0.022 | [] | 0.6682 |
| *solitary* | -2794.83 | 0.011 | 0.048 | 0.0001 |
| *subsocial* | -2802.16 | 0.022 | [] | 0.9989 |
| *parasocial* | -2799.76 | 0.024 | 0.000 | 0.0282 |
| *primitively eusocial* | -2800.99 | 0.020 | [] | 0.1257 |
| *advanced eusocial* | -2800.07 | 0.025 | 0.005 | 0.0408 |
| *Apidae* | -2801.8 | 0.022 | [] | 0.3926 |
| *Halictidae* | -2801.94 | 0.023 | [] | 0.5078 |
| *Xylocopinae* | -2801.2 | 0.023 | [] | 0.1660 |
| *null* | -2802.16 | | | |

| | | | | |
|---|---|---|---|---|
| *IRS* | | | | |
| *cluster A* | -15553.09 | 0.093 | [] | 0.0800 |
| *cluster B* | -15551.99 | 0.093 | 0.133 | 0.0217 |
| *cluster C* | -15554.59 | 0.098 | [] | 0.8023 |
| *cluster D* | -15554.31 | 0.096 | [] | 0.4243 |
| ***E. tridentata*** | **-15540.88** | **0.093** | **0.328** | **0.0000*** |
| *solitary* | -15553.59 | 0.094 | [] | 0.1498 |
| *subsocial* | -15554.15 | 0.097 | [] | 0.3286 |
| *parasocial* | -15553.54 | 0.095 | [] | 0.1414 |
| *primitively eusocial* | -15554.55 | 0.098 | [] | 0.7046 |
| *advanced eusocial* | -15548.04 | 0.091 | 0.150 | 0.0003 |
| *Apidae* | -15554.10 | 0.098 | [] | 0.3049 |
| *Halictidae* | -15551.45 | 0.095 | 0.161 | 0.0117 |
| *Xylocopinae* | -15554.32 | 0.098 | [] | 0.4375 |
| *null* | -15554.63 | | | |
| *inR-2* | | | | |
| *cluster A* | -11081.62 | 0.044 | [] | 0.7063 |
| *cluster B* | -11081.47 | 0.044 | [] | 0.5106 |
| ***cluster C*** | **-11066.22** | **0.049** | **0.007** | **0.0000*** |
| *cluster D* | -11072.73 | 0.050 | 0.024 | 0.0000 |
| *E. tridentata* | -11070.30 | 0.041 | 0.100 | 0.0000 |
| *solitary* | -11081.14 | 0.046 | [] | 0.2937 |
| *subsocial* | -11079.98 | 0.043 | [] | 0.0643 |
| *parasocial* | -11079.98 | 0.043 | [] | 0.0643 |
| ***primitively eusocial*** | **-11066.22** | **0.049** | **0.007** | **0.0000** |
| *advanced eusocial* | -11081.69 | 0.045 | [] | 0.9924 |
| *Apidae* | -11081.17 | 0.046 | [] | 0.3092 |
| *Halictidae* | -11079.28 | 0.043 | 0.085 | 0.0283 |
| *Xylocopinae* | -11080.43 | 0.044 | [] | 0.1126 |
| *null* | -11081.69 | | | |
| *InR* | | | | |
| *cluster A* | -9219.15 | 0.032 | [] | 0.5627 |
| *cluster B* | -9219.32 | 0.031 | [] | 0.9414 |
| *cluster C* | -9214.99 | 0.033 | 0.011 | 0.0032 |
| *cluster D* | -9219.03 | 0.032 | [] | 0.4422 |
| ***E. tridentata*** | **-9208.16** | **0.028** | **0.082** | **0.0000** |
| *solitary* | -9219.04 | 0.032 | [] | 0.4533 |
| *subsocial* | -9219.32 | 0.031 | [] | 0.9587 |
| *parasocial* | -9219.32 | 0.031 | [] | 0.9414 |
| *primitively eusocial* | -9214.99 | 0.033 | 0.011 | 0.0032 |
| *advanced eusocial* | -9215.23 | 0.028 | 0.045 | 0.0042 |
| *Apidae* | -9216.15 | 0.033 | 0.016 | 0.0118 |
| *Halictidae* | -9216.71 | 0.030 | 0.095 | 0.0224 |
| *Xylocopinae* | -9219.21 | 0.032 | [] | 0.6419 |
| *null* | -9219.32 | | | |

**Table S6.** Results for linear regression of dN/dS values of six focal genes with PCs 1-4.

| | **PC1** | | **PC2** | | **PC3** | | **PC4** | |
|---|---|---|---|---|---|---|---|---|
| | $R^2$ | *p* | $R^2$ | *p* | $R^2$ | *p* | $R^2$ | *p* |
| *ISR* | 0.00 | 0.81 | 0.00 | 0.88 | 0.00 | 0.83 | 0.05 | 0.35 |
| *InR* | 0.07 | 0.45 | 0.31 | 0.09 | 0.00 | 0.82 | 0.08 | 0.43 |
| *InR-2* | 0.06 | 0.51 | 0.13 | 0.33 | 0.28 | 0.14 | 0.00 | 0.89 |
| *TOR* | 0.28 | 0.07 | 0.00 | 1.00 | 0.24 | 0.11 | 0.08 | 0.37 |
| *dunce* | 0.02 | 0.56 | 0.13 | 0.18 | 0.18 | 0.09 | **0.34** | **0.01** |
| *syx1a* | **0.32** | **0.02** | 0.03 | 0.52 | 0.07 | 0.34 | 0.00 | 0.93 |
